# EEG decodability of facial expressions and their stereoscopic depth cues in immersive virtual reality

**DOI:** 10.1101/2025.08.18.670974

**Authors:** Felix Klotzsche, Ammara Nasim, Simon M. Hofmann, Arno Villringer, Vadim Nikulin, Werner Sommer, Michael Gaebler

**Affiliations:** Max Planck Institute for Human Cognitive and Brain Sciences, Department of Neurology, Leipzig, Germany; Humboldt-Universität zu Berlin, Department of Psychology, Germany; Carl von Ossietzky Universität Oldenburg, Germany; Fraunhofer Institute for Telecommunications, Heinrich Hertz Institute (HHI), Department of Artificial Intelligence, Berlin, Germany; Department of Physics and Life Science Imaging Center, Hong Kong Baptist University, Hong Kong, China; Faculty of Education, National University of Malaysia, Kuala Lumpur, Malaysia

**Keywords:** Face perception, depth perception, EEG, immersive virtual reality, emotional expressions, multivariate decoding, eye movements

## Abstract

Face perception typically occurs in three-dimensional space, where stereoscopic depth cues enrich the perception of facial features. Yet, most neurophysiological research on face processing relies on two-dimensional displays, potentially overlooking the role of stereoscopic depth information. Here, we combine immersive virtual reality (VR), electroencephalography (EEG), and eye tracking to examine the neural representation of faces under controlled manipulations of stereoscopic depth. Thirty-four participants viewed computer-generated faces with neutral, happy, angry, and surprised expressions in frontal view under monoscopic and stereoscopic viewing conditions. Using time-resolved multivariate decoding, we show that EEG signals in immersive VR conditions can reliably differentiate facial expressions. Stereoscopic depth cues elicited a distinct and decodable neural signature, confirming the sensitivity of our approach to depth-related processing. Yet, expression decoding remained robust across depth conditions, indicating that under controlled frontal viewing, the neural representation of behaviorally distinguishable facial expressions is invariant to binocular depth cues. Eye tracking showed that expression-related gaze patterns contained comparable information but did not account for neural representations, while depth information was absent in gaze patterns—consistent with dissociable representational processes. Our findings demonstrate the feasibility of EEG-based neural decoding in fully immersive VR as a tool for investigating face perception in naturalistic settings and provide new evidence for the stability of expression representations across depth variations in three-dimensional viewing conditions.

## Introduction

Faces are among the most relevant visual input for humans. They play a crucial role in social communication, enabling us to recognize and distinguish individuals, as well as to infer detailed semantic information about them, such as their emotional states (Bruce & Young, 1986; Jack & Schyns, 2015). Humans are able to detect subtle differences between faces and facial expressions, although they are composed of relatively few key features (e.g., one mouth, one nose, two eyes) and share a similar configuration. Moreover, faces and the emotions they convey are processed and evaluated more rapidly than other objects of comparable visual complexity (Willis & Todorov, 2006). The sophisticated processing of facial stimuli is supported by a hierarchical cortical network, ranging from retinotopic areas in the early visual cortex, which process single features, to face-specific visual areas like the fusiform face area (FFA) that encode view-invariant, abstract representations of faces (Calder & Young, 2005; Duchaine & Yovel, 2015; Freiwald et al., 2016; Haxby et al., 2000).

In the real world, humans constantly process faces in complex, three-dimensional (3D) settings. However, laboratory studies of face perception usually simplify these experiences, relying on two-dimensional (2D) stimuli presented on flat screens. Such conditions omit key features of natural vision—for example, binocular depth information. Binocular disparity is a visual feature which is automatically and (seemingly) effortlessly extracted from the retinal inputs and integrated into a percept with stereoscopic depth: Each eye captures a slightly different image, which the cerebral cortex integrates—together with further visual cues such as relative size, blur, shading, perspective, texture gradients, or motion as well as with sensory signals from other modalities—to gain depth information about the shape of an object or the configuration of a scene (Parker, 2007; Welchman, 2016). During this fusion into a stereoscopic representation, coarse percepts change and refine over time (Menz & Freeman, 2003). Among the various available depth cues, binocular information plays a special role for the subjective experience of stereoscopic depth (Barry, 2010; Vishwanath, 2014). However, binocular disparity information is neither necessary (Vishwanath & Hibbard, 2013) nor sufficient (Cumming & Parker, 1997) for a subjective impression of stereoscopic depth. The precise contribution of binocular disparity information for the brain’s processing of complex stimuli like faces—and, in particular, how it interacts with the representation of other visual features of the same object—is still unclear.

As humans often encounter faces at close range—where binocular disparity provides important cues (Cutting & Vishton, 1995)—depth information may play a key role in face processing. Indeed, stereoscopic depth modulates the recognition and representation of faces and facial expressions: It improves participants’ ability to recognize the same person from different viewpoint angles or distances (Burke et al., 2007; C. H. Liu & Ward, 2006; H. Liu et al., 2020). C. H. Liu et al. (2006) observed impaired performance when recognition conditions differed from encoding conditions in the availability of spatial depth cues (stereoscopic vs monoscopic), suggesting that neural encoding depends on whether faces are processed with or without stereoscopic depth information. Hakala et al. (2015) demonstrated that stereoscopically presented facial expressions evoke stronger emotional responses than their monoscopic counterparts, with natural depth cues amplifying negative reactions to angry faces. Children (3–6 years old) are faster in recognizing emotional facial expressions in stereoscopic as compared to monoscopic conditions (Wang et al., 2017). Collectively, these findings suggest that stereoscopic depth cues can shape natural face processing, highlighting the importance of binocular information in how humans perceive and respond to faces and their expressions.

Here, we used electroencephalography (EEG) in combination with immersive virtual reality (VR) and eye tracking to investigate how the human brain processes faces and facial expressions in the presence of stereoscopic depth cues. Due to its high temporal resolution, EEG is a powerful tool for studying the temporal dynamics of face perception (for reviews see Brunet, 2023; Rossion, 2014). Previous EEG research has identified several characteristic components of the event-related potential (ERP), each emerging within a specific time window and reflecting a distinct processing stage of cortical face processing: The earliest component associated with face processing is the P1 component (80–120 ms) which has been linked to low-level visual analysis of faces (Rossion & Caharel, 2011). It is followed by the N170 (130–200 ms), generated in the fusiform gyrus (Gao et al., 2019), which encodes structural face recognition and differentiates facial expressions and identities (Rossion & Jacques, 2012). Around 250–300 ms after stimulus onset, the Early Posterior Negativity (EPN) indexes reflexive attentional orienting toward emotional contents and its selection for further processing (Schindler & Bublatzky, 2020). Finally, the Late Positive Complex (LPC; 400–600 ms) reflects higher-order cognitive evaluation of faces and other stimuli of motivational relevance (Schacht & Sommer, 2009; Schupp et al., 2006).

These well-established ERP components provide informative temporal markers of the sequential stages of face processing in the human brain. However, because they are typically derived by averaging over predefined electrode clusters, they may overlook more subtle, spatially distributed patterns of neural activity. Multivariate decoding—used to probe whether specific stimulus information is present in high-dimensional neurophysiological signals (e.g., Kamitani & Tong, 2005; for a recent review, see Peelen & Downing, 2023)—extends this approach by exploiting the full spatiotemporal structure of the EEG data and optimally weighting channels on the level of the individual participant (Carrasco et al., 2024; Grootswagers et al., 2017). By doing so, it can offer greater sensitivity, detecting stimulus-specific information embedded in distributed activity patterns—information that might not be apparent in classical ERP analyses.

In the following, we refer to such decodable information as *representation*. While this usage is controversial (see Fallon et al., 2023, for an overview), it aligns with how the term is commonly used by many neuroscientists—where *representation* is often approximated with *decodability* and reflects mutual information between a pattern of brain activity and an experimental variable of interest (Hebart & Baker, 2018).^1^ Neuroscientists use decodability as a proxy for determining *where* and *when* information about a feature is available in the brain (Grootswagers et al., 2017). EEG-based decoding offers high temporal resolution, enabling researchers to track the sequential stages at which such information emerges. Quantifying the contribution of individual EEG channels to decoding performance permits a coarse spatial interpretation of the underlying signals. Subsequent source reconstruction can map these patterns to their cortical generators, thereby identifying the brain regions most relevant for the representation of the decoded feature.

In the present study, we applied this approach to investigate facial expression representations while selectively manipulating the presence of stereoscopic depth cues. To achieve this, we used immersive VR with head mounted displays (HMDs) to deliver binocular stimulation. Immersive VR is increasingly adopted by cognitive and perceptual scientists—not only for its ability to present stereoscopic depth (Choi et al., 2023; Draschkow, 2022; Tarr & Warren, 2002; Thurley, 2022) but also because the computer-generated stimulation of the (almost) entire visual field enables high levels of experimental control and facilitates various opportunities for stimulus modification. Building on these developments, cognitive neuroscientists have begun combining immersive VR with EEG to investigate neural processing of face stimuli under 3D viewing conditions (Nolte et al., 2024; Sagehorn et al., 2023; Sagehorn, Johnsdorf, et al., 2024; Sagehorn, Kisker, et al., 2024; Schubring et al., 2020).

However, combining HMDs with EEG recordings presents several challenges. Placing the HMD over an EEG cap introduces additional sources of noise, potentially obscuring signals of interest (Tauscher et al., 2019; Weber et al., 2021). Larger stimulus eccentricities due to the wider field of view compared to conventional computer screens can affect visually evoked ERP components (Klotzsche et al., 2023). In addition, while conventional laboratory studies often restrict eye movements to control retinotopic stimulus presentations, VR paradigms typically permit free exploration of the environment, which leads to eye movement-related artifacts and confounds in the EEG data and may require different analytical approaches (Nolte et al., 2024, 2025). Finally, little is known about how the mere addition of binocular depth alters the neural processing of relevant stimuli. Overall, it remains unclear to what extent findings from conventional 2D screen-setups generalize to 3D settings that include stereoscopic depth information.

The present study had two main objectives. First, we addressed a methodological challenge: whether time-resolved EEG decoding can recover facial expression information in an immersive VR setup, where signal quality and noise sources differ from conventional laboratory settings. Second, we examined whether the neural representations of facial expressions are modulated by the presence of stereoscopic depth cues—examining both whether depth information alters expression-related activation patterns and whether stereoscopic depth itself is a decodable feature in the EEG. To contextualize these findings, we performed complementary source reconstruction and eye tracking analyses to characterize neural and eye movement-related origins of both facial expression and stereoscopic depth cue representations.

**Figure 1:**
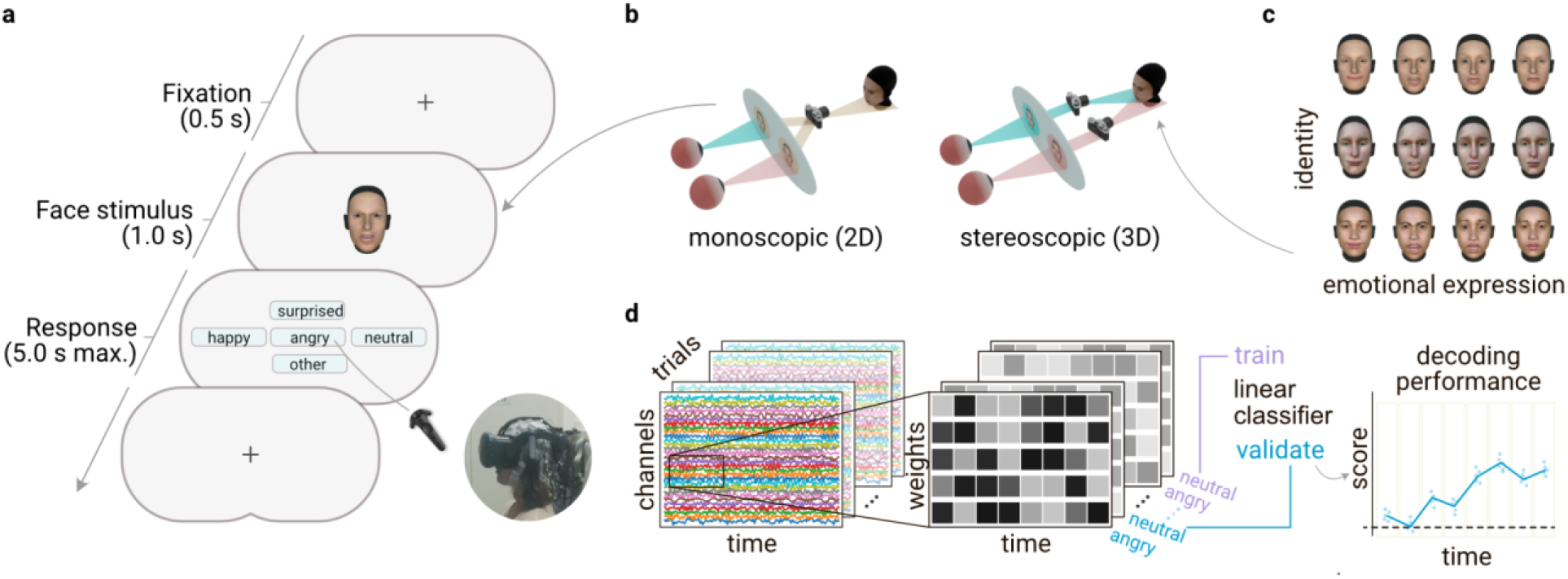
Experimental paradigm and decoding analysis. **(a)** In each trial of the emotion recognition task, participants fixated centrally before a face stimulus was presented for 1,000 ms. Subsequently, they selected the recognized emotional expression using the VR controller. **(b)** Face stimuli were presented either with (right) or without (left) stereoscopic depth information. For the stereoscopic condition, a 3D model of the face was rendered using two slightly offset virtual cameras (a “binocular VR camera rig”), providing binocular depth cues. For the monoscopic condition, the same 3D model was first rendered from a single virtual camera to produce a flat 2D texture, eliminating stereoscopic disparity. This texture was then presented to both eyes via the binocular VR camera, removing binocular depth cues while maintaining all other visual properties. **(c)** Overview of all stimulus combinations: three different face identities (rows) expressing four different emotional facial expressions (columns: happy, angry, surprised, neutral). Face identity was irrelevant for the emotion recognition task. Face models were originally created and evaluated by Gilbert et al. (2021). **(d)** Schematic of the decoding approach: A multivariate linear classifier (logistic regression) was trained on successive time windows of EEG data, treating each channel as a feature. The emotional expression shown in the respective trial served as the classification label. Using a 5-fold cross-validation, the classifier was trained repeatedly on 80% of the trials and its predictions were validated on the remaining 20%. This procedure yielded decoding performance scores per time window, reflecting the amount of available stimulus information in the EEG data at the corresponding time in the epoch.

## Results

### EEG Decoding

#### Decodability of facial expressions

To assess whether EEG activity contained information about the viewed facial expression, we trained time-resolved linear classifiers separately for each participant on multichannel EEG data and evaluated their cross-validated performance at successive time points after stimulus onset. Decoding performance was quantified as ROC-AUC, with values above 0.5 indicating above-chance classification. We tested for significance using cluster-corrected permutation tests. Facial expression information was reliably decodable from the EEG data, with decoding performance significantly exceeding chance-level (Figure 2). For multiclass decoding (distinguishing between the four facial expressions: neutral, angry, happy, and surprised), we observed a large interval with above-chance decoding performance extending from 94 ms after stimulus onset until the onset of the response panel (1,000 ms after stimulus onset). The highest decoding performance (ROC-AUC; average of individual maxima: M = 0.61, SD = 0.03, 95% CI [0.60, 0.62]) was observed for most participants at a median (Mdn) delay of 414 ms (SD = 315.81, 95% CI [306.25, 521.75]).

**Figure 2:**
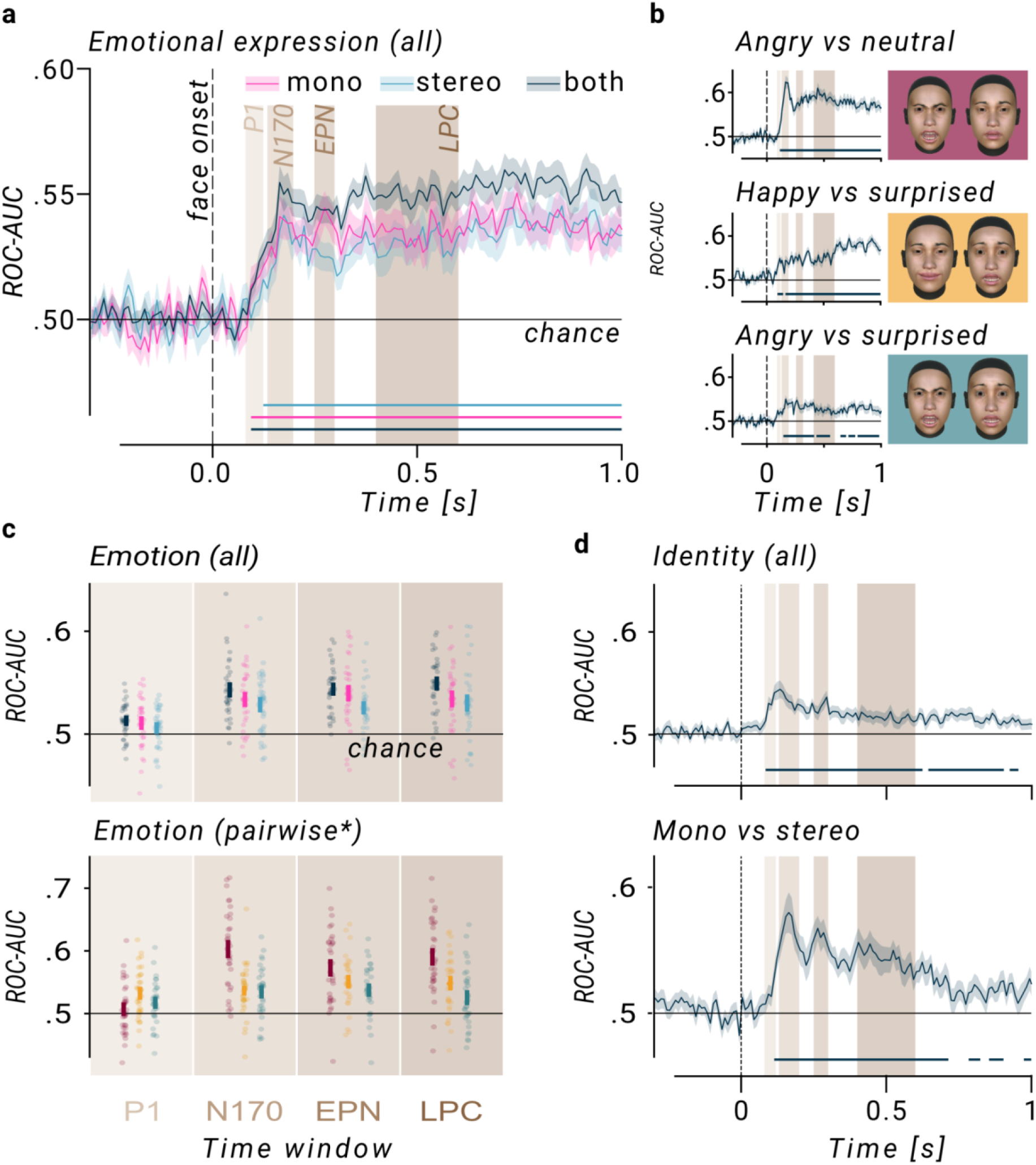
Time-resolved classification performance for different decoding targets and conditions. **(a)** Classification performance (mean ± 1 SEM) of the decoder distinguishing between the four facial expressions, trained separately for the mono- and stereoscopic viewing conditions, as well as on data pooled across both depth conditions. Horizontal lines at the bottom indicate clusters where decoding was significantly above chance. Shaded rectangles in the background mark relevant ERP time windows associated with face processing. **(b)** Selection of three (out of six) binary contrasts which underlie the multiclass classification in (a). *Angry vs neutral*: Highest decoding performance. *Happy vs surprised*: classification performance lacks an early peak and only rises later in the trial. *Angry vs surprised*: Lowest (but still significant) decoding performance. **(c)** Average performance per time window, depth condition (*top*; colors as in (a)), and binary contrast (*bottom*; *: selection and colors as in (b)). Thick bars: mean ± 1 SEM. **(d)** Time-resolved classification performance (mean ± 1 SEM) for the task-irrelevant decoding targets: identity of the stimulus face (*top*) and presence vs absence of stereoscopic depth information (*bottom*). ROC-AUC: area under the receiver operating characteristic.

To determine which expression pairs contributed to the multiclass effect, we additionally trained binary classifiers for all six pairwise contrasts between emotional expressions. We then summarized decoding performance within four canonical face-processing time windows (P1, N170, EPN, and LPC) to characterize the temporal profile of expression-related information. Figure 2, Supplementary Figure S3, and Supplementary Table ST1 show the decoding results for these binary decoders. For each contrast, we found at least one significant cluster, demonstrating above chance-level decoding also for all binary classifiers. Descriptively, different emotion pairs exhibited distinct pairwise-decoding profiles, varying in average decoding performances and temporal dynamics (e.g., the contrast between angry and neutral emotions showed higher and earlier average decodability compared to the contrast between angry and surprised). We further found a significant main effect of the time window on the ROC-AUC (F(3,96) = 22.01, p < .001) which was driven by a significantly lower decoding performance in the P1-window compared to all other time windows. The remaining time windows did not differ significantly for the multiclass decoding approach. For the binary classifiers (contrasting pairs of facial expressions), we observed again a significant main effect of time window (F(3,96) = 21.13, p < .001), a significant main effect of contrast (F(5,160) = 9.43, p < .001), as well as a significant interaction (F(15,480) = 9.02, p < .001). The P1-window yielded the lowest decoding performance across all binary decoders. Only for one out of the six contrasts (“surprised-vs-happy”), the decoding performance was significantly above chance during this time window. All other binary contrasts did not differ in their average ROC-AUC and were not significantly above chance level in the P1 window. In the remaining time windows, all contrasts, except for “angry-vs-surprised” in the LPC window, performed significantly above chance level. The decoders distinguishing “angry-vs-neutral” and “angry-vs-happy” showed significantly higher decoding performance than the other binary classifiers. This difference was most pronounced in the N170 and the LPC time windows. Details are depicted in Figure 2 and Supplementary Figure S4.

Together, these analyses show that facial expression information can be reliably decoded from EEG while observing faces in an immersive VR headset. Temporal and contrast-specific differences indicate that decodability depends on both the time window and the specific pair of expressions being distinguished.

#### The role of stereoscopic depth cues

To test whether stereoscopic depth cues modulated the neural representations of facial expressions, we repeated the expression decoding analyses separately for monoscopic and stereoscopic trials and compared decoding performance between the viewing conditions. The multiclass decoders validated on trials with and without stereoscopic depth information differed neither in terms of the maximal decoding performance (mono: M = 0.61, SD = 0.02, 95% CI [0.60, 0.62]; stereo: M = 0.61, SD = 0.03, 95% CI [0.60, 0.62]; t(32) = −0.49, p = .627) nor for the latency of the peak decoding performance (mono: Mdn = 384.00 ms, SD = 287.71, 95% CI [285.83, 482.17]; stereo: Mdn = 524.00 ms, SD = 386.06, 95% CI [392.28, 655.72]; t(32) = −1.04, p = .305). This was confirmed by a cluster-corrected comparison of the decoding performance at each time-point which did not indicate a significant difference between the two depth conditions. In a four-by-two rmANOVA which modeled the average decoding performance as a function of the time window (four levels: P1, N170, EPN, LPC) and the depth condition (two levels: mono- and stereoscopic), only time window was a significant predictor (F(3,96) = 17.82, p < .001), but neither depth condition (F(1,32) = 1.55, p = .222) nor its interaction with time window (F(3,96) = 1.00, p = .395). Also for all the binary classifiers (distinguishing pairs of emotions), we observed no significant differences between the two depth conditions, neither for peak decoding performance nor for its latency (Table ST1). This absence of depth-related modulation was further supported by cross-decoding analyses: classifiers trained only on trials from one of the viewing conditions (with or without stereoscopic depth information) performed equally well when tested on trials from the same or the other depth condition (see Supplementary Figure S5). Thus, adding stereoscopic depth cues did not measurably enhance, reduce, or alter the profile of EEG-based facial expression decoding, suggesting that the decodable expression information was largely invariant to the presence of binocular disparity cues.

To scrutinize further whether any information about the presence or absence of stereoscopic depth cues was present in the EEG data, we made the depth condition itself the decoding target of a binary classifier (i.e., decoding whether the data came from a trial in the mono- or the stereoscopic condition). Figure 2d shows the time-resolved decoding performance of this decoder. Above-chance decodability was indicated by a long interval of significant decoding performance starting 114 ms after stimulus onset (end: 714 ms) and multiple shorter intervals at later time points in the epoch. On average, a peak decoding score of M = 0.65 (SD = 0.07, 95% CI [0.62, 0.67]) was observed for most participants at Mdn = 184 ms after stimulus onset. Descriptively, the decoding performance peaked early (peak decoding time: Mdn = 184 ms, SD = 204.17, 95% CI [114.34, 253.66]) and continuously decreased toward the end of the epoch. We observed a similar temporal profile of the decoding performance when classifying the identity of the stimulus face (rows in Figure 1c)—another task-irrelevant stimulus feature—from the EEG data (peak decoding score M = 0.60, SD = 0.03, 95% CI [0.59, 0.61]; peak decoding time: Mdn = 134 ms, SD = 244.62, 95% CI [50.54, 217.46]; Figure 2d). Thus, stereoscopic depth information was clearly present in the EEG signal, even though it did not modulate expression decoding. Its decoding profile resembled that of face identity, consistent with depth acting as a task-irrelevant stimulus feature in the present task.

#### Source reconstruction

To characterize which sensor-level patterns contributed to decoding, we transformed classifier weights into spatial activation patterns (Haufe et al., 2014) and projected these patterns onto a standard cortical surface using eLORETA source reconstruction. We summarized the most informative cortical regions within the same canonical time windows used for the decoding analyses. Figure 3 depicts these decoder-related patterns and a reconstruction of the associated cortical sources. Classification of task-relevant features (facial expression) as well as task-irrelevant features (identity, depth condition) primarily involved occipital, parietal, and temporal sources. Early after stimulus onset (P1, N170, EPN windows), the most informative sources for facial expression classification were located in the primary and early visual cortex. At later stages (LPC window), sources in the parietal cortex became more prominent contributors. Supplementary Table ST2 lists the most relevant sources for the binary classifiers, which distinguish between pairs of facial expressions. Supplementary Figure S9 visualizes the associated cortical source reconstructions. For the contrasts with the highest decoding performance (“angry-vs-neutral” and “angry-vs-happy”), primary and early visual cortices remained the most informative sources, even during the LPC time window. In contrast, the other pairwise comparisons between two emotional expressions showed more variable results, with regions along both the ventral and dorsal processing streams playing dominant roles.

**Figure 3:**
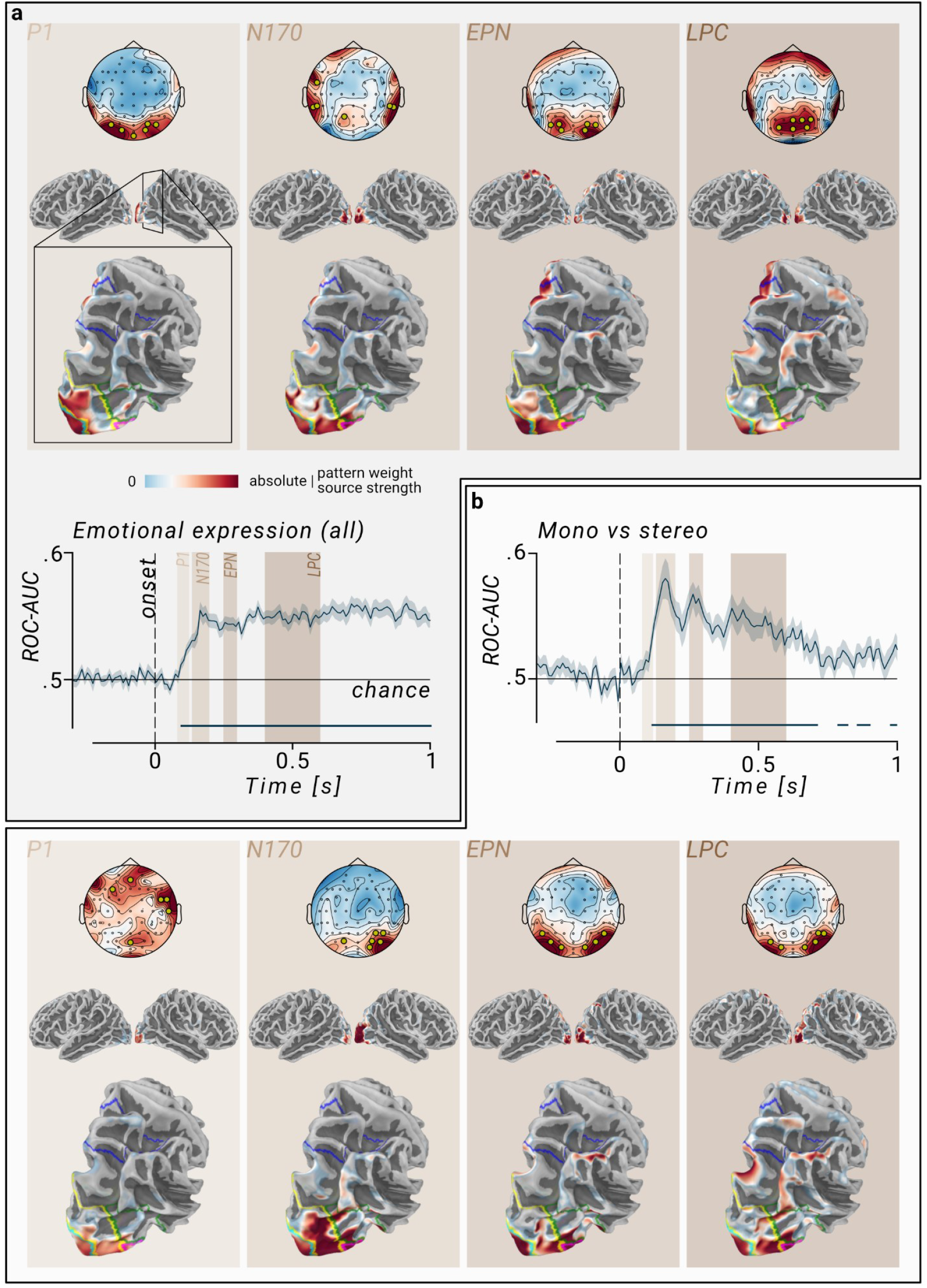
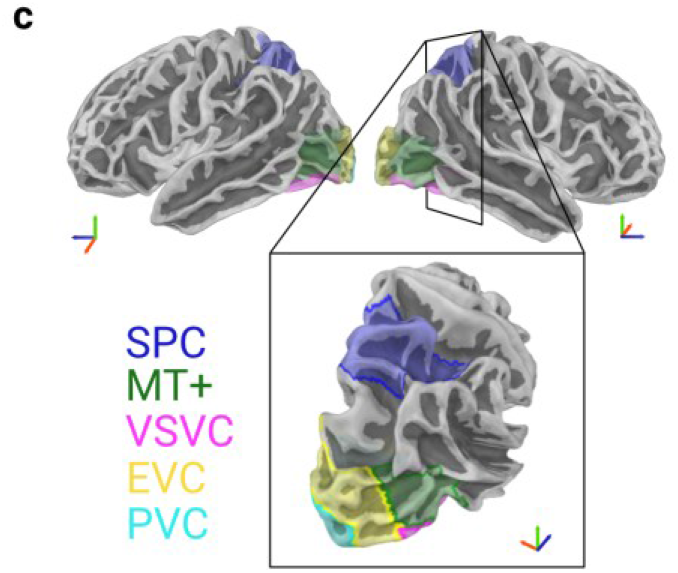
Spatial patterns and localized sources related to decoding (a) facial expressions and (b) the availability of stereoscopic depth information. **(a)** *Top*: Classification of the four facial expressions. Spatial patterns of the classifier (top row) and their projection onto the cortical surface using eLORETA (second and third row) for the four time windows. Colors indicate the absolute, normalized weight of each sensor or cortical parcel (warmer colors represent higher absolute weights). In the topographies, the six most informative channels (top 10%) are highlighted in yellow. Projections on the cortical surface are masked to show only the top 5% most informative parcels. Second row: Lateral view on both hemispheres. Third row: View on occipital and parietal cortices of the right hemisphere. Colored lines on the cortex mark the outlines of relevant regions (see (c)). *Bottom*: Time course of decoding performance for multiclass classification of the four emotional expressions (as in Figure 2). **(b)** As in (a) [in reversed order], but for the classifier trained to distinguish between trials with and without stereoscopic depth information. **(c)** Outlines of relevant cortical regions, following the reduced parcellation atlas by Glasser et al. (2016) and Mills (2016). *SPC: Superior Parietal Cortex; MT+: MT+ Complex and Neighboring Visual Areas; VSVC: Ventral Stream Visual Cortex; EVC: Early Visual Cortex; PVC: Primary Visual Cortex (V1)*.

The most informative sources for the classifier trained to distinguish trials with and without stereoscopic depth information were located along the ventral visual processing stream and in the MT+ complex (see Figure 3 and Supplementary Figure S8). Similarly, the decoding of face identity relied primarily on signals originating from the ventral visual processing stream during the N170 time window (Supplementary Figure S8). In contrast, during the two later time windows (EPN and LPC), identity decoding shifted to rely most on information from parietal regions (see Supplementary Table ST2 for an overview).

Overall, the source estimates primarily implicated occipital visual cortical regions with later parietal contributions depending on the decoded feature.

### Eye Tracking

#### Saccade and fixation patterns

To check if the dynamics of the EEG decoding performance could reflect eye movements, we analyzed the concurrently recorded eye tracking data (combined gaze between left and right eye) for a subsample of participants (n=17), for whom eye tracking data were available. We tested whether gaze behavior varied as a function of facial expression, stereoscopic depth condition, and time window. Figure 4b shows the average fixation heatmaps on the face stimuli separately for the different facial expressions. Supplementary Figure S6 shows the corresponding heatmaps separately for each of the four time windows. Participants mostly kept fixation at the center of the face. Towards later time windows (after about 200 ms), there was a notable shift of fixations towards the lower half of the face. This observation was supported by a 4×2×4 MANOVA which modeled the average fixation position as a function of facial expression, depth condition, and time window. The multivariate result was significant for the predictor time window (Wilk’s lambda = 0.93, F(6, 1022) = 6.14, p < .001) but not for the other predictors or any interaction. Univariate follow-up analyses (rmANOVA and post-hoc t-tests) demonstrated that vertical fixation locations differed significantly between the LPC time window and all other time windows. No significant differences were observed among the remaining time windows, and averaged horizontal fixation positions did not differ significantly across any time window.

**Figure 4:**
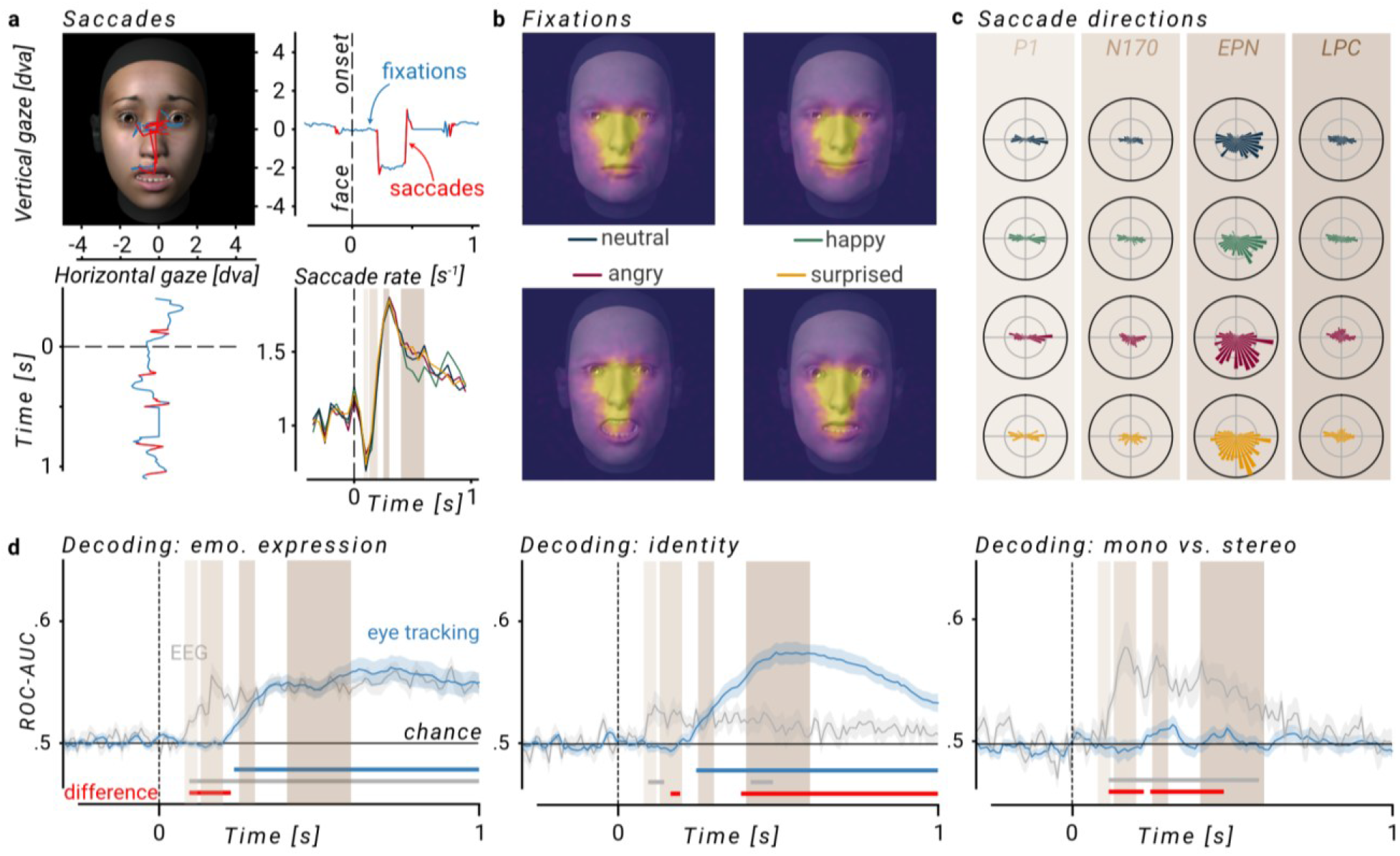
Eye tracking analyses and comparison with the EEG decoding results. **(a)** *Top left*: Exemplary gaze trace (one trial) on a stimulus face. Saccades are plotted in red, fixations in blue. The fixation cross preceding the stimulus was displayed at (0,0). *Top right*: Vertical component of the same gaze trace over time. *Bottom left*: Horizontal gaze component over time. *Bottom right*: Saccade rate (saccades per second) over the time of the trial across all trials and participants. Colors like in (b) and (c). We observed a dip in saccade rate right after stimulus onset, followed by a sharp increase peaking during the EPN time window. **(b)** Heatmaps showing fixation distributions within the first second after stimulus onset, across all trials and participants, separated by emotional expression. **(c)** Circular histograms of saccade directions (polar angles relative to the preceding fixation), plotted per time window and emotional expression (colors like in (b)). Saccade counts are normalized by the length of the time window. Most saccades occurred during the time window of the EPN (an ERP component associated with reflexive attention to the stimulus), predominantly in downward (especially for angry and surprised faces) or lateral directions. **(d)** Decoding performance of classifiers trained on the gaze data (spherical coordinates) for different decoding targets. EEG-based decoding performance is overlaid in gray for comparison (in the subset of participants with eye tracking data). Horizontal bars at the bottom of each plot indicate time points with decoding significantly above chance level. Red bars mark significant differences in performance between eye tracking and EEG-based decoding (two-sided, cluster-corrected *t*-test). Note, EEG decoding results shown here are based on the subsample with eye tracking data (n=17), resulting in lower scores compared to Figures 2 and 3.

We next quantified whether saccade frequency, amplitude, and direction showed systematic condition-dependent changes that could parallel the EEG decoding dynamics. In a generalized linear model (GLM; Poisson, log-linear) modeling the number of saccades (Pseudo R² = 0.90, Supplementary Table ST3) only time window was a significant predictor (Wald χ²(3) = 115.81, p < .001). Neither facial expression (Wald χ²(3) = 2.42, p = .491) nor depth condition (Wald χ²(1) = 1.47, p = .225) or any interactions explained significant amounts of variance. The average saccade amplitude did not differ significantly between facial expressions, depth conditions, or time windows (all p-values > .150). Furthermore, the model did not explain a substantial portion of the variance in the data (Gaussian, log-linear GLM; Pseudo R² = 0.03). Figure 4c shows the distribution of saccade directions as a function of the time window and expression. A MANOVA which modeled the average saccade direction (two-dimensional, spherical coordinates), indicated a significant main effect of time window (Wilk’s lambda = 0.96, F(6, 968) = 3.22, p = .004) and a significant interaction between time window and emotional expression (Wilk’s lambda = 0.94, F(18, 968) = 1.65, p = .04), but no significant effects for the other predictors.

Thus, gaze behavior changed over the course of stimulus viewing, with later fixations shifting toward the lower face and saccade patterns varying primarily as a function of time. However, these effects were only weakly expression-dependent and showed no systematic modulation by stereoscopic depth condition.

#### Decodability from gaze data

To increase the sensitivity of our eye tracking analyses, we used multivariate decoding to test whether gaze data contained information about the facial expressions. As for the EEG data, we trained a set of time-resolved, linear classifiers to test the decodability of relevant attributes—however, now from the two-dimensional gaze data (i.e., the orientation of the combined eyes in spherical coordinates). Using cluster-corrected permutation testing, we demonstrated significant decoding performance for the classifier distinguishing between the four emotional expressions. This was indicated by a cluster of significant decoding scores which started 230 ms after stimulus onset and extended to the end of the epoch (1,000 ms). The highest decoding performance (ROC-AUC; M = 0.59, SD = 0.03, 95% CI [0.57, 0.60]) was observed for most participants at Mdn = 700 ms (SD = 230.04, 95% CI [590.65, 809.35]). Using the four time windows which we also used to bin the EEG data (P1, N170, EPN, LPC), we found a significant effect of the time window on the decoding performance (F(3,48) = 26.78, p < .001), with significant pairwise differences between all time windows apart from P1 vs N170 and EPN vs LPC (in other words, no differences between the two early and between the two late time windows). Only for the two later time windows, EPN and LPC, the performance of the decoder was significantly above chance level.

As for the EEG analyses, we additionally examined all six pairwise expression contrasts to determine whether gaze-based decodability differed between specific emotion pairs. We found significant decoding performance for all binary classifiers (pairwise distinction of facial expressions) supported by cluster-corrected permutation testing against chance-level. Further, a rmANOVA comparing the average decoding performance across the binary contrasts and the four time windows revealed significant main effects for the decoding contrast (F(5,80) = 3.12, p = .013) and the time window (F(3,48) = 29.70, p < .001), but no significant interaction (F(15,240) = 1.50, p = .106). Post-hoc tests (pairwise t-tests, Bonferroni corrected) showed significant differences in the decoding performance between all time windows, apart from P1 and N170 which did not differ significantly. Later time windows yielded higher decoding performances than early time windows. The post-hoc pairwise comparisons of the contrasts did not reveal any significant differences (after Bonferroni correction).

Thus, facial expression information was also present in gaze behavior but emerged primarily in later time windows. As in the EEG analyses, gaze-based decodability varied across time windows and showed contrast-dependent differences.

#### Stereoscopic information

As for the EEG data, to test whether stereoscopic depth affected gaze-based expression information, we repeated the gaze-decoding analyses separately for monoscopic and stereoscopic trials and compared decoding performance between the two viewing conditions. Gaze-based expression decoding did not differ between monoscopic and stereoscopic viewing conditions. For the decoding of the facial expressions, there were no significant differences in peak performance (mono: M = 0.60, SD = 0.03, 95% CI [0.58, 0.61]; stereo: M = 0.59, SD = 0.03, 95% CI [0.57, 0.60]; t(16) = 1.88, p = .079) or timing (mono: Mdn = 741.67 ms, SD = 268.15, 95% CI [614.20, 869.14]; stereo: Mdn = 650.00 ms, SD = 301.43, 95% CI [506.71, 793.29]; t(16) = 1.11, p = .283). This finding was indirectly supported by a non-significant cluster-based permutation test contrasting the time-resolved performances for expression decoding in the mono- and the stereoscopic condition. Similarly, no binary classifier showed significant performance differences between the two depth conditions.

We then asked whether depth condition itself, as well as face identity, could be decoded from gaze direction. In contrast to the EEG data, a separate classifier based on the time-resolved eye tracking data could not differentiate between the two depth conditions (monoscopic vs stereoscopic; see Figure 4d). However, a classifier trained to differentiate between the face identities achieved above chance-level decoding, indicated by a cluster of time-points with significant decoding accuracy, beginning at 242 ms after stimulus onset and continuing until the onset of the response panel (peak decoding score: M = 0.60, SD = 0.03, 95% CI [0.58, 0.61]; peak decoding time: Mdn = 550.00 ms, SD = 231.29, 95% CI [440.05, 659.95]; Figure 4d).

Similar to the EEG data, gaze behavior carried information about facial expression and face identity. Gaze-based expression decoding was not modulated by stereoscopic depth. However, in contrast to the EEG, information about the presence of binocular depth cues was not decodable from gaze data.

#### The relationship between EEG and eye tracking based decoding performances

Finally, to assess whether EEG-based decoding could be explained by gaze behavior, we directly compared the decoding profiles obtained from EEG and eye tracking data in the same subset of participants. We examined both participant-level associations in decoding strength and latency, and time-resolved differences between modalities. On the group level, there was no significant correlation between the performance of the decoder trained to distinguish facial expressions based on EEG data and the one based on eye tracking data. This applied to peak decoding performance (r(15) = 0.059, p = .821), latency of the peak performance (r(15) = −0.160, p = .540), and average decoding performance across the observation window (r(15) = −0.031, p = .907). Decoding performance based on EEG data surpassed decoding based on eye tracking data early in the trial, as indicated by a significant cluster between 94 and 224 ms (see Figure 4d, left). For the remaining time of the observation window, there was no significant difference between EEG and eye tracking based decoding of facial expressions. By contrast, the decoders for facial *identity* displayed a different relationship between their performance profiles (see Figure 4d, middle): the EEG-based decoding again ramped up earlier than the eye tracking-based decoding (first significant cluster: 164–194 ms), but at later stages, eye tracking-based decoding outperformed EEG-based decoding (second significant cluster with opposite-signed cluster mass: 384–1,000 ms). In decoding the depth condition, the classifier trained on EEG data outperformed the classifier trained on eye tracking data (significant clusters: 114–224 ms and 244–474 ms), with the latter never reaching a performance level above chance.

Overall, the comparison between modalities suggests that EEG- and gaze-based decoding captured partly overlapping but dissociable sources of stimulus processing.

### Behavior

To verify that participants correctly recognized the displayed expressions, we analyzed their categorical responses in an emotion recognition task. Recognition accuracy was computed by comparing participants’ responses with the intended expression labels based on the FACS-validated stimulus set (i.e., the FACS ratings provided by Gilbert et al., 2021). The results are shown in the form of confusion matrices in Supplementary Figure S1. Recognition rates differed across the emotional expressions (F(3,96) = 6.81, *p* < .001) but were high for all expressions (correct: M = 90.73%, SD = 16.98, 95% CI [89.06, 92.41]). The lowest recognition level was found for neutral faces (correct: M = 80.32%, SD = 25.12, 95% CI [71.75, 88.89]) which were classified second most frequently as “other” (confusion rate: M = 17.39%, SD = 23.51, 95% CI [9.37, 25.41]). We did not observe a difference in recognition rate for the mono-(M = 90.96 %, SD = 16.79, 95% CI [88.10, 93.82]) and the stereoscopic depth conditions (M = 90.51 %, SD = 17.32, 95% CI [87.55, 93.46]). Accordingly, there was neither a significant main effect of depth condition (F(1, 32) = 1.85, p = 0.183) nor a significant interaction with the emotional expression (F(3, 96) = 1.37, p = .260). Thus, participants reliably recognized the intended facial expressions, and recognition performance was not measurably affected by the presence of stereoscopic depth cues.

We additionally assessed whether the stimuli differed in perceived emotional intensity, and whether stereoscopic depth altered these subjective intensity ratings. At the end of the experiment, we asked participants to rate for each of the 24 stimuli (4 emotions, 3 face identities, 2 depth conditions; Figure 1c) the intensity of the displayed emotional facial expression using the 9-point SAM scale for arousal (Supplementary Figure S2). We observed a significant main effect of emotion (F(3,93) = 118.95, p < .001): Angry faces received the highest intensity ratings (M = 7.82, SD = 0.93, 95% CI [7.50, 8.14]), while neutral faces received the lowest (M = 2.69, SD = 1.66, 95% CI [2.11, 3.26]). Happy (M = 4.60, SD = 1.47, 95% CI [4.09, 5.11]) and surprised faces (M = 6.15, SD = 0.96, 95% CI [5.81, 6.48]) fell in between. Neither the main effect of the depth condition (F(1,31) = 0.01, p = .932) nor its interaction with the emotional expression was significant (F(3,93) = 0.34, p = .794). Overall, subjective intensity ratings differed strongly between emotional expressions, but not between monoscopic and stereoscopic presentations.

## Discussion

We investigated the neurocognitive processes underlying the recognition of facial expressions in immersive VR and if the presence of stereoscopic depth cues changes the neural representations of face stimuli. In each trial, we showed a computer-generated face, which displayed one out of four facial expressions. As the main measure of interest we used time-resolved, multivariate decoding from EEG data to infer how much information about the stimulus is present in the brain signals. Our main decoding target was the emotional expression shown by the face—which the participants had to indicate at the end of each trial. Facial expressions could be decoded from EEG data with robust above-chance accuracy. At the same time, the decoding performance was not influenced by the availability of stereoscopic depth information. We demonstrate that information about the presence of stereoscopic depth cues is present in the EEG signals nevertheless: With a separate classifier we could successfully distinguish mono-from stereoscopic trials. The temporal profile of this decoding differed from decoding the expressions but resembled the temporal profile of decoding the identity of the stimulus face, another task-irrelevant feature. We therefore conclude that in neurocognitive processing of face stimuli, stereoscopic depth information is treated like other task-irrelevant stimulus features and—at least as long as the behavioral task can be easily solved without integrating this information—it does not modify the (measurable) neurophysiological representation.

### Successful decoding of emotional expression

Our decoding results reveal that the EEG data recorded in our VR setup contained information about the emotional expression of the stimulus faces. Previous studies have demonstrated that similar linear EEG classifiers could distinguish between different facial expressions presented as grayscale, two-dimensional images on conventional computer screens (Smith & Smith, 2019). Our results replicate and extend these findings by showing that time-resolved multivariate decoding yields comparable results when using naturalistic, 3D face stimuli in immersive VR settings. Such setups are gaining interest in human perception and cognition research for their ability to provide more naturalistic stimulation—closer to how people perceive and experience the world—thereby potentially improving the generalizability of experimental findings to real-world contexts (Cisek & Green, 2024). Yet, VR also introduces substantial challenges compared to the traditional, more restrictive setups. For example, combining EEG with a VR headset can lower the signal-to-noise ratio by introducing artifacts from mechanical pressure on the electrodes, muscular activity caused by head movements or headset weight, or electrical noise from the device and its cables (Tauscher et al., 2019; Weber et al., 2021). Despite these challenges, our study shows that EEG signal quality in controlled VR settings is sufficient to reliably discriminate between facial expressions and identities by using linear classifiers applied to an observer’s EEG data. In this context, multivariate decoding is especially valuable, condensing high-dimensional EEG data into a one-dimensional, time-resolved, and interpretable metric—decoding performance—via optimized, data-driven weighting of EEG channels. Cross-validation ensures that the classification results are generalizable and not due to overfitting. In contrast, traditional ERP analyses rely on strong a priori assumptions (e.g., preselected electrodes) and often require extensive post-hoc corrections for multiple comparisons, yet rarely employ cross-validation. Prior studies have shown that multivariate decoding offers higher sensitivity than conventional ERP analyses (Carrasco et al., 2024; Grootswagers et al., 2017), which is particularly relevant for data prone to low signal-to-noise ratios. Future VR studies on face processing in immersive contexts may build on our findings to inform their methodological choices and anticipate effect sizes.

Consistent with the findings by Smith & Smith (2019), our EEG decoding results showed distinct patterns for task-relevant and task-irrelevant features: emotional expression (task-relevant) yielded sustained decodability after it surpassed chance level, whereas the decoding performance for task-irrelevant features (here: face identity and availability of stereoscopic depth cues) peaked early, followed by temporal decline (Figure 2).

Systematically controlling for low-level visual features (e.g., spatial frequencies, luminance) of naturalistic face stimuli presented in immersive VR is challenging. Because we did not independently control or manipulate these features, our analyses cannot determine their contribution to expression-or identity-related decoding. We therefore do not claim that our observations exclusively reflect high-level face processing. Indeed, some of our results suggest that the decoding of facial expressions may have relied on low-level visual differences between the stimulus classes: the highest decoding performance was achieved for facial expression pairs with particularly salient visual differences (e.g., the visibility of the teeth). By contrast, our central manipulation of stereoscopic depth used the same face models and facial expressions in both viewing conditions, preserving low-level visual features while only varying the availability of binocular disparity.

### The role of stereoscopic depth cues

We hypothesized that 3D viewing—by more closely approximating natural perceptual contexts—would elicit more robust and distinct neural representations, thereby enhancing decoding performance. Contrary to this prediction, we found no significant differences in decoding performance between the two conditions for either a task-relevant (expression) or a task-irrelevant (identity) decoding target. Stereoscopic information did not enhance decoding performance, alter its temporal dynamics, or affect the spatial distribution of the classifier weights. These findings were corroborated by cross-condition generalization tests (cross-decoding), where classifiers trained and tested on stereoscopic trials performed equally well when tested on monoscopic trials, and vice versa (Supplementary Figure S5). If 3D and 2D percepts relied on fundamentally different neural activation patterns, we would expect reduced generalization across conditions. However, our results provide no evidence for such a difference, suggesting that the neurophysiological activation patterns underlying facial expression categorization remain largely consistent regardless of the presence of stereoscopic depth information.

If the subjective percepts in the two conditions are markedly different, why do the underlying neural activation patterns associated with the decoded features show no significant differences? Importantly, our main analyses employed activation patterns optimized for decoding specific target variables (facial expression, face identity), not representing the brain’s holistic response to the stimuli. Within this scope, 3D and 2D conditions did not show differences. However, when directly decoding the presence of stereoscopic depth cues, we identified significant distinctions in the neural responses, peaking around 180 ms after stimulus onset and subsequently declining—similar to the temporal profile for decoding face identity, another task-irrelevant feature (Figure 2d). We thereby demonstrate that the distinction between viewing conditions with and without stereoscopic depth cues was detectable in our neurophysiological data. This adds to previous findings regarding binocular depth processing gained with different methods (from single cell recordings in non-human primates to fMRI in humans; for an overview see Welchman, 2016) and stimuli (from face pictures; Chou et al., 2021; to abstract random dot motion; Parker, 2007). Our study addresses a relevant research gap by demonstrating that EEG data in a VR setup with naturalistic, computer-generated stimuli can track stereoscopic depth processing.

Time-resolved decoding profiles also provide insights into the temporal organization of face and depth processing. Expression decoding, once significantly above chance-level, remained sustained throughout the trial, whereas depth decoding peaked around 180 ms after stimulus onset and then declined. Sustained expression decoding may reflect the task-relevance of emotional information, which engages ongoing evaluative processing (Schacht & Sommer, 2009). In contrast, the peak and subsequent decline of depth decoding performance—resembling the decoding profile of face identity, another task-irrelevant feature (Smith & Smith, 2019)—suggest that stereoscopic depth was processed as a perceptual feature during initial face perception, but progressively less prominently thereafter. The peak of depth decoding falls within the N170 time window, previously associated with structural face encoding (Rossion & Jacques, 2012). Therefore, binocular disparity information might get integrated into the face processing stream at this point. However, the absence of depth-related modulation of expression decoding across all time windows suggests that depth and expression information were processed in parallel, without significant interference—at least under conditions where the behavioral task does not require depth information.

Concretely, our findings indicate disparity-related responses that distinguish mono- and stereoscopic viewing conditions without detectably altering the activation patterns underlying facial expression decoding. This finding may reflect the respective decoders exploiting information from different cortical areas.

### Cortical sources

In an exploratory analysis, we therefore applied source reconstruction to project the spatial patterns associated with the decoders back onto the cortical surface. For the classification of emotional expressions, primary and early visual cortices contributed most substantially, especially shortly after stimulus onset (P1, N170, and EPN windows). This dominance was particularly evident in the binary contrasts with the highest decoding performance (“angry-vs-neutral”, “angry-vs-happy”; see Supplementary Figure S9 and Supplementary Table ST2). In contrast, lower-performing binary classifiers (e.g., “angry-vs-surprised”) relied primarily on regions in lateral occipital and posterior inferotemporal cortex. This suggests that decoding was more reliable when information about the contrast was also present in primary or early visual cortices. This aligns with previous studies which found that early visual cortices, which mostly process low-level stimulus features, support effective visual input decoding (Kamitani & Tong, 2005; Lützow Holm et al., 2024; Wilson et al., 2024). Accordingly, the contrasts with the highest decoding performance expressed the most salient visual differences between the involved faces. For example, angry faces were characterized by a widely opened mouth showing both rows of teeth, whereas neutral and happy faces had a closed mouth with no teeth visible. The decoding of these contrasts may have relied heavily on these low-level feature representations. Nevertheless, contrasts with more subtle low-level visual differences still yielded above chance-level decoding, primarily drawing on brain regions higher in the visual processing hierarchy. When two facial expressions lacked sufficient differences in their low-level visual features, decoding appeared to rely more on signals from cortical areas known to process abstract features of facial expressions. Overall, our findings indicate that also for naturalistic face stimuli the decoding-based approach does not simply exploit arbitrary features, rely solely on low-level visual features, or exclusively depend on abstract representations of the observed facial expression.

The decoding of the (task-irrelevant) face identities was mostly informed by areas in the ventral visual stream—in particular during the N170 time window where the decoding performance was highest. This aligns with previous findings that the N170 reflects activity of the fusiform face area (Gao et al., 2019) and neighbouring ventral areas which process information relevant for distinguishing different faces (Eimer, 2011; Grill-Spector et al., 2017; Kanwisher & Yovel, 2006). Running the source reconstruction separately for the mono- and stereoscopic conditions yielded only slightly different distributions across the cortical surfaces as compared to running it on the pooled data (see Supplementary Table ST2). We attribute these variations to the fact that splitting the data set leads to less robust source estimates and therefore a higher variance. Overall, we did not observe a systematic difference between the distribution of sources in the mono-as compared to the stereoscopic viewing condition, congruent with the numeric decoding results.

The decoder trained to explicitly categorize the depth condition (i.e., separating mono-from stereoscopic trials) mostly relied on information from areas in the ventral visual stream and the MT+ complex. This is consistent with previous findings that stereoscopic depth is processed across multiple stages of the visual stream, in particular in dorsal and ventral visual areas (Preston et al., 2008). Compared with the cortical areas primarily associated with facial expression decoding in our study, this result supports the claim that depth decoding may have drawn on partly different cortical sources, potentially explaining why depth was decodable without detectably altering facial expression decoding. Related evidence comes from Deligiannis et al. (2025) who found that binocular disparity of face stimuli modulates fMRI responses in depth-sensitive regions but has a very limited effect on the activity in face-sensitive areas.

Our spatially resolved source reconstruction has limitations: We used a common average head model without individual anatomical brain scans, relied on the canonical locations in the 10-20 system without participant-specific registration, and faced potential spatial distortions (i.e., electrode displacement) from the VR headset worn over the EEG cap. Despite these constraints, we successfully recorded and analyzed spatially interpretable signals that meaningfully map onto the cortical surface.

### Computer-generated faces as stimuli

We used computer-generated stimuli, unlike most EEG studies on human face processing which have typically relied on standardized photographs (e.g., from the Chicago Face Database; Ma et al., 2015). Computer-generated stimuli can be precisely tailored to the specific requirements of each study whereas photographs are less modifiable and challenging to alter without introducing visible distortions. For example, to produce gradual (i.e., parametrically adjustable) versions of facial expressions, “morphing” between versions of two existing images has been used (Steyvers, 1999). In recent years, more complex generative algorithms have gained momentum which allow for the creation of artificial but increasingly naturalistic face stimuli both in 2D and 3D (e.g., Barthel et al., 2025; Feng et al., 2021). Here, we used 3D models and predefined blend shape modifications to generate a set of static facial expressions. These expressions had been evaluated in 2D (i.e., rendering 2D images of the generated 3D models) by an independent sample of both experts and non-experts (Gilbert et al., 2021), ensuring participants could clearly distinguish them—increasing the likelihood of obtaining separable EEG signals. However, the very pronounced expressions and their somewhat mask-like appearance, resulting from the use of blend shapes, admittedly limits their representativeness of real-world faces. Our results establish that robust EEG decoding is achievable in immersive VR when using clearly distinguishable face stimuli, serving as a foundation for future studies using more ambiguous and naturalistic facial stimuli to explore how these affect both behavioral responses and EEG decodability.

### Behavioral recognition

Participants were clearly able to recognize the displayed emotional facial expressions. We designed the stimuli by selecting facial expressions (i.e., FACS configurations) with empirically validated high recognition rates (Gilbert et al., 2021), minimizing ambiguities in decoding arising from subjective indecisiveness. This approach ensured that potential challenges encountered during neural decoding could be ascribed to a lack of information in the EEG signal itself (e.g., low signal-to-noise ratio), rather than to the relevant information not being encoded in the first place. While future studies using more ambiguous facial expressions may reveal additional aspects of stereoscopic information processing, here we sought to minimize the risk of uninterpretable null results due to non-significant decoding scores. A prior EEG decoding study using clearly distinguishable facial expressions achieved only modest above-chance performance (Smith & Smith, 2019), and VR-EEG setups are likely to exacerbate noise-related reductions in effect size. We therefore prioritized maximizing decoding feasibility in VR-EEG as a foundational step for subsequent work, with a special focus on the introduction of stereoscopic depth cues.

Building on our results and effect sizes, future studies can now more confidently work their way towards more ambiguous facial expressions while studying the concurrent neurophysiology. We cannot exclude the possibility that for more ambiguous facial expressions, for which decoding is more difficult (both for the human observer and the EEG decoder), stereoscopic depth cues become more critical. Similarly, in computer vision-based recognition of human facial expressions, it has been shown that, particularly for low-intensity expressions, models incorporating 3D information outperform those using only 2D information (Savran et al., 2012).

### Emotional intensity of the stimuli

Participants’ subjective ratings at the end of the experiment indicated differences in emotional intensity between the facial expressions and the different identities. Stereoscopic depth had no influence on the intensity ratings.

While the stimulus intensity may partially explain the EEG decoding results, for example, through arousal contagion (Herrando & Constantinides, 2021), this alone cannot fully account for the results. While emotional arousal has widespread effects in the human brain which can be read out by similar decoding approaches (Hofmann et al., 2021), these are typically not localized in early visual cortices—one of the most informative brain areas in the present study. Furthermore, face identity was decodable with similar accuracy to expressions despite substantially weaker differences between the identities in terms of emotional intensity. Finally, stereoscopic viewing conditions were successfully differentiated based on the EEG, despite no differences in the emotional intensity ratings between with and without stereoscopic depth cues.

However, the above argumentation may hold less for later times in the trial (>500 ms after stimulus onset), during which the decoding performance declined for the task-irrelevant decoding targets (identity and depth condition) while it was constant for the decoding of the facial expressions. Also, some of the most informative sources shifted from early visual cortex to parietal and ventral temporal regions, which have been found to be modulated by emotional arousal (Greene et al., 2014; Lettieri et al., 2019; Wade-Bohleber et al., 2020). This suggests that early decoding reflects visual feature processing, while later stages may incorporate emotional arousal effects.

### Eye tracking results and their relation to the EEG results

The EEG decoding might have been influenced by eye movements (Mostert et al., 2018), as participants could freely explore the faces (for 1,000 ms) after initial central fixation. Analyses of fixation locations and saccade parameters revealed that the saccade rate peaked around the EPN time window, with later fixations targeting the lower face—consistent with previous findings (Schurgin et al., 2014). Saccade direction, especially in the later part of the trial, was modulated by the facial expression—also consistent with previous findings (Calvo et al., 2018; Schurgin et al., 2014; Vaidya et al., 2014). Time-resolved linear classifiers showed comparable decoding performances from eye tracking and EEG data (Figure 4)—with similar temporal profiles—raising the question of whether EEG decoding might, in the end, simply reflect eye movements rather than neurophysiological representations of stimulus features (see Mostert et al., 2018, for a more detailed discussion of this argument and potential underlying mechanisms).

Importantly, multiple analyses indicate that eye movements do not fully account for the observed EEG patterns. First, if the decodable activity in the EEG data was directly caused by eye movements (e.g., due to eyeball motion or by retinal image shifts), both time series should be perfectly time-aligned or show a slight lead for eye tracking signals. Instead, we observed a different pattern: eye tracking decodability consistently lagged behind EEG signals by 100–200 ms (see Figure 4d). This delay cannot be attributed to temporal imprecision during data recording. To compensate for potential lag in eye tracking data recorded with the HTC Vive Pro Eye, we post-hoc aligned it with the EEG time series, by cross-correlating the eye tracking data with the electrooculogram channels. We identified and corrected a similar delay as reported by Stein et al. (2021). Stimulus information was therefore decodable from EEG *before* the concurrent eye tracking data, making it implausible that eye movements caused the patterns we observed in the EEG. Second, if EEG decodability were merely a byproduct of the eye movements, decoding scores from both modalities should be correlated (e.g., participants with highly/late decodable eye movements should exhibit high/late EEG decoding peaks). Contrary to this assumption, we found no systematic relationship between EEG and eye tracking-based decoding metrics on the participant level (see Supplementary Figure S7). Finally, while the decoding performance curves for EEG and eye tracking look similar when classifying facial expressions, the patterns differ for the task-irrelevant decoding targets (Figure 4d). Most prominently, the depth condition—while well decodable from EEG—was entirely undecodable from eye tracking data. It should be noted that eye tracking data were only available for 17 of the 33 participants included in the EEG analyses, which limits the generalizability of these conclusions. The dissociations, despite being based only on a subsample, make it unlikely that our EEG decoding results merely reflect eye movements. While the similarity in some dynamics suggests shared information between the modalities, the dissociations strongly indicate that EEG decoding provides complementary insights, capturing aspects of neurocognitive processing beyond what eye tracking data alone can reveal—for example, the presence or absence of stereoscopic depth information.

## Conclusion

In sum, our results demonstrate that emotional facial expressions can be reliably decoded from EEG in immersive VR, with no clear advantage observed under conditions involving stereoscopic depth cues. While the presence of stereoscopic depth elicited distinct neural signatures, it did not affect the decodability of other, orthogonal face features—whether task-relevant or irrelevant. This suggests that, in controlled facial emotion recognition tasks using clearly distinguishable, static stimuli, the brain treats stereoscopic depth like other independent stimulus features—at least from the perspective of decodable EEG representations. Our findings establish a methodological foundation for EEG-based decoding in immersive VR and invite future research to explore how richer, more ambiguous, or dynamic face stimuli—potentially viewed from different perspectives—might leverage stereoscopic depth information in behaviorally and neurophysiologically meaningful ways.

## Methods

### Participants

We acquired data from 34 healthy, young, female adults (age: M = 26.65, SD = 4.56, 5 left-handed). Persons wearing glasses could not participate in the study to facilitate compatibility with the eye tracker. We ensured intact stereoscopic vision in all participants using a Titmus test (*Fly-S Stereo Acuity Test*, Vision Assessment Corporation, Hamburg, Germany). The study was approved by the ethics committee of the Department of Psychology at the Humboldt-Universität zu Berlin and participants provided their written consent prior to participation. Participants were compensated with 12 € per hour.

The final sample consisted of data from 33 participants for the EEG analyses (incomplete data for 1 participant) and 17 participants for the eye tracking control analyses (due to technical failure, no eye tracking data were stored for the remaining 16 participants).

### Setup

We recorded participants’ electroencephalogram (EEG) and electrooculogram (EOG) using a LiveAmp amplifier (BrainProducts GmbH, Gilching) and 60 active electrodes according to the 10-20 system in an electrode cap (actiCAP snap; BrainProducts). Four EOG electrodes were placed below the eyes and next to the outer canthi. All electrodes were referenced to electrode FCz (ground: FPz). EEG and EOG were sampled at a rate of 500 Hz and using a hardware-based low-pass filter at 131 Hz (third-order sinc filter, −3 dB cutoff). We ensured that at the beginning of the experiment, the impedances of all electrodes were below 25 kΩ.

During the VR part of the study, participants were seated and wore a VR headset (HTC Vive Pro Eye, HTC, Taiwan) with an integrated eye tracker (sampling rate: 120 Hz). We adjusted the headset at the beginning of the experiment to match the interpupillary distance of the participant. The VR headset was positioned on top of the EEG cap (covered by a disposable shower cap). The flat design of the actiCAP snap electrodes, combined with the cushioning of the HTC Vive Pro’s back section, facilitated an even weight distribution, minimizing excessive pressure on any individual electrode. A custom facial interface cushion with recesses at designated locations prevented pressure on the frontal EEG electrodes (Fp1/2).

### Software

For the implementation of the experiment, we used the Unity game engine (v2020.3.3f1; Unity Technologies) in combination with SteamVR (Valve Corporation) and ran it on a VR-ready PC (Intel Core i9-9900K, 3.6 GHz, 32 GB RAM, NVIDIA RTX 2080Ti GPU, Windows 10). The computer was connected to the EEG amplifier via an analog port (D-SUB 25) to enable synchronization of the EEG data with the experimental events. We used the EDIA toolbox (Klotzsche et al., 2025)—an extension of the Unity Experiment Framework (v2.3.4; Brookes, 2017/2019; Brookes et al., 2020)—for structuring the experiment, recording events, and tracking the behavior (incl. eye movements). The eye tracker was interfaced using the SRanipal SDK and EDIA to allow for sampling the eye tracking data at 120 Hz. To synchronize the data streams (i.e., stimulus onset events, eye tracking, EEG), we used custom C# scripts to send analog triggers (EEG) and store associated timestamps (eye tracking and event data). We recorded EEG and EOG data using the BrainVision Recorder software (v1.22.0101; BrainProducts). Before and after the VR-based part of the experiment, participants filled in questionnaires implemented in SoSciSurvey (Leiner, 2019). Data analysis was performed using Python (v3.10.4), Scikit-learn (v1.5.1; Pedregosa et al., 2011), SciPy (v1.14.0; Virtanen et al., 2020), statsmodels (v0.14.2; Seabold & Perktold, 2010), and NumPy (v1.26.4; Harris et al., 2020).

### Stimuli

The stimulus set entailed computer-generated faces from three digital humans, each showing four different facial expressions (angry, happy, neutral, surprised; see Figure 1c), yielding 12 static face stimuli in total. We generated the faces using the open-source software makehuman and the FACSHuman plugin (Gilbert et al., 2021), based on material provided by the first author of the plugin. Gilbert et al. (2021) provide validations of 2D, grayscale versions of these stimuli. Two independent raters, professionally trained in the Facial Action Coding System (FACS; Ekman & Friesen, 1978), had approved the adherence of the emotional expressions to the FACS criteria and their recognizability was demonstrated in a sample of naïve participants in an online experiment (Gilbert et al., 2021). For the present study, we selected the four emotional expressions which were best recognized and distinguished from one another in this validation experiment. We chose only female faces and we manually masked the hair and the ears of the 3D models (using the Blender software; v2.93, Blender Development Team, 2021) to decrease task-unrelated differences between the stimuli. Each face stimulus measured approximately 24 cm in vertical height (from the crown of the head to the base of the neck), roughly matching the dimensions of an adult human face in physical reality. We presented the faces against a gray background and at a distance of 1.37 virtual meters from the perspective of the participant, so that for the observer each face spanned approximately 10 degrees of visual angle (dva) along its vertical axis. Stimuli were presented centrally in the visual field in a way that the virtual head frontally faced the participant irrespective of the orientation of the participant’s head (i.e., in *local space*). Head movements, therefore, did not lead to displacements of the stimuli within the visual field. For the 3D condition, we displayed the regular three-dimensional model of the face stimuli, yielding a stereoscopic view in the VR headset. The VR software renders two slightly different viewpoints of the scene to the displays in the headset, thereby enabling stereoscopic depth cues. In the 2D condition, we replaced the 3D model by a two-dimensional “portrait shot” taken by a single cyclopean (virtual) camera and displayed as a 2D plane (canvas) in the Unity scene. The 2D image was size- and position-aligned with the (invisible) 3D head model and then rendered to the displays in the headset, yielding an impression very similar to the rendering in the 3D condition, just without stereoscopic depth information (Figure 1b). The percept in the 2D condition was comparable to viewing a life-size portrait picture from a distance of 1.37 m.

### Task

In the main task of the experiment, participants performed an emotion recognition task (Figure 1a). In each trial, after a fixation period of 500 ms, we displayed for 1,000 ms a single face stimulus centrally in the visual field of the participant. After the offset of the face, the participant was presented with five response buttons arranged in a cross-like fashion centrally in the visual field (see Figure 1a). The buttons displayed the four facial expression categories (neutral, angry, happy, surprised) as well as the option “other”. Using a VR controller, participants indicated their response by pointing a ray toward the corresponding virtual button. The assignment of the buttons to the positions on the cross was randomized across trials. Participants could therefore only decide where to press once the buttons were visible, thus avoiding preparatory activity during the presentation of the face stimulus. The next trial started immediately after the response or automatically 5,000 ms after stimulus offset.

In 50% of the trials, the face stimuli were shown with stereoscopic depth information. In the remaining trials, the 2D version of the face was presented. The experimental manipulations (facial expression, identity, depth condition) were fully interleaved and pseudorandomized, ensuring a uniform distribution of all experimental conditions within each block. Participants completed six experimental blocks with 120 trials each, yielding 720 trials in total. After each block, the experiment was paused and participants could take a short break (incl. removal of the VR headset and re-calibration of the eye tracker).

Before the start of the first block, participants completed 24 training trials to get familiar with the task and the stimuli. In these trials, each combination of face identity, facial expression, and depth condition was shown once in a random sequence. Data from the training trials were not analyzed. After the main block of the experiment, participants performed an intensity rating of all face stimuli. To this end, all 24 combinations of face identity, facial expression, and depth condition were shown in a random sequence in the VR headset. This time, the face stimulus was visible permanently (until the participant had responded) above a 9-point self-assessment manikin (SAM) scale (Bradley & Lang, 1994) that allowed the participant to indicate the intensity (i.e., the degree of emotional arousal) of the displayed facial expression (see Supplementary Figure S2). Participants selected the according level with the VR controller. No EEG or eye tracking data were recorded for this part.

### EEG preprocessing

We processed the EEG data with MNE-Python (v1.2.3; Gramfort et al., 2013). First, we identified and rejected channels which were noisy throughout the experiment based on their power spectrum and visual inspection. Using ICA decomposition (extended Infomax), we then removed artifacts caused by blinks and eye movements. For fitting the ICA solution, we used a separate copy of the data which we filtered between 1 and 40 Hz (FIR filter with a Hamming window of length 1651 samples, lower/upper passband edge: 1.00/40.00 Hz, lower/upper transition bandwidth: 1.00/10.00 Hz, lower/upper −6 dB cutoff frequency: 0.50/45.00 Hz). From this copy, we extracted epochs of 1,300 ms in length, starting 300 ms before and ending 1,000 ms after the onset of the face stimuli. Epochs with particularly noisy EEG signals were identified using the autoreject software (v0.4.3; Jas et al., 2017) and removed before fitting the ICA weights. Based on their correlation with the bipolar EOG channels and visual inspection, we identified components related to blinks and other eye movements. Using only the remaining ICA weights (i.e., setting the weights of affected components to zero), we cleaned a separate version of the data. This dataset was filtered between 0.1 and 40 Hz (FIR filter with Hamming window of length 16,501 samples, lower/upper passband edge: 0.10/40.00 Hz, lower/upper transition bandwidth: 0.10/10.00 Hz, lower/upper −6 dB cutoff frequency: 0.05/45.00 Hz) to preserve slow ERP components. Again, we extracted epochs of 1,300 ms in length, starting 300 ms before and ending 1,000 ms after the onset of the face stimuli. For baseline-correction, we subtracted the mean voltage during a 200 ms baseline interval (i.e., the 200 ms before stimulus onset) from each epoch. Subsequently, we used the autoreject package for local (i.e., per participant, sensor, and epoch) interpolation and trial rejection.

### EEG decoding

We trained and cross-validated time-resolved linear classifiers on the EEG data to test the decodability of the facial expressions from the participant’s brain activity. A separate decoding model was validated for each participant using all available EEG channels. The samples used for training and testing the classifier were formed by averaging mini-batches of three trials from the same class. Using averaged data from mini-batches has been found to improve the performance of classifiers trained on EEG data due to suppression of noise (Adam et al., 2020; Grootswagers et al., 2017). Along the time-dimension, we downsampled the signal using a moving average of five samples (10 ms) with no overlap. For each time point, we then trained a logistic regression model (solver: liblinear, L2-regularization: λ = 1.0) on the data from all 60 EEG sensors and assessed its decoding performance by means of a stratified 5-fold cross-validation. In each fold, the model was trained on 80% of the mini-batches and tested on the remaining 20% while we kept the class proportions equal between train and test sets. Comparing the model predictions with the ground truth (FACS ratings of the facial expressions) in the test set, we calculated the area under the receiver operating characteristic (ROC-AUC) to assess the model’s decoding performance. To obtain a robust estimate, we performed this decoding procedure 50 times for each participant and time point—varying the random allocation of trials into mini-batches and randomly splitting samples into train and test sets. An overall decoding score per time point and participant was obtained by averaging the ROC-AUC scores across all repetitions.

We conducted cluster-based permutation tests on the group-level to determine whether the decoding performance was significantly above chance-level during the first second after stimulus onset. To this end, we calculated a one-sided paired t-test for each time point, testing if the actual decoding score was significantly higher than chance (ROC-AUC = 0.5). The cluster-based permutation procedure allowed for a correction of the number of tests (i.e., the number of time points). For a physiological interpretation of the decoding results, we calculated the *patterns* of the classifier (by multiplying the covariance of the EEG data with the filter weights; Haufe et al., 2014) for each time point.

To test the decodability of the facial expression (i.e., a four-class classification problem), we trained the above-mentioned model by applying a *one-vs-rest* (i.e., one vs the three remaining conditions) multiclass extension of the binary classifier. We further evaluated separate models for each two-sided contrast (i.e., six pairwise contrasts between the four emotional expressions) for enhanced interpretability.

We calculated the mean decoding score within four distinct time windows, based on canonical ERP components associated with face expression processing (P1, N170, EPN, LPC), to facilitate the interpretability of the decoding results across time. Using repeated-measures analyses of variance (rmANOVAs) and post-hoc *t-*tests (Bonferroni corrected), we modeled the decoding performance as a function of the *time window* (four levels) and the binary decoding *contrast* (six levels).

To test whether the presence of stereoscopic depth cues impacts the decodability, we performed the same decoding procedure separately with data from the mono- and stereoscopic depth conditions. We then compared the resulting decoding performances at each time window using two-sided, paired *t*-tests with cluster-correction. We also compared the monoscopic and the stereoscopic depth condition in terms of the peak decoding performance (highest decoding performance across all time points) and its latency using paired, two-sided *t*-tests. Further, we used a rmANOVA with the factors *time window* (P1, N170, EPN, LPC) and *depth condition* (mono-vs stereoscopic) to model the decoding scores. Finally, we used the same decoding procedure to make the *depth condition* itself the decoding target (i.e., predicting if a sample was measured under mono- or stereoscopic viewing conditions), as well as the *identity* of the stimulus face (three classes; see Figure 1c)—both task-irrelevant features of the stimulus.

### Source reconstruction

We applied source reconstruction to project the spatial patterns obtained from the decoding process onto the cortical surface. Specifically, we used exact low-resolution tomography analysis (eLORETA; Pascual-Marqui, 2007) to localize the sources corresponding to the extracted components. The reconstruction utilized the standard average forward model from FreeSurfer (Fischl et al., 1999) as provided by MNE-Python, restricting the solution to dipoles with fixed orientations perpendicular to the cortical surface. Using eLORETA, we constructed spatial filters for each voxel from the leadfield matrix. The sources were then computed by multiplying the demixing matrix with the spatial patterns (Haufe et al., 2014) derived from the classifiers trained in EEG channel space. For multiclass classifiers, this procedure was performed separately for each underlying binary (one-vs-rest) classifier, and the normalized source time courses were averaged (per time point) to produce a single overall source time course for the multiclass decoder. To determine the most informative source within the canonical ERP time windows, we calculated the average weight of each vertex over the respective time intervals, identified the vertex with the highest average weight, and classified its corresponding brain region using the reduced parcellation atlas (23 labels per hemisphere) provided by Glasser et al. (2016) and Mills (2016).

### Eye tracking analyses

We analyzed the eye tracking data (combined gaze data for left and right eye as provided by the eye tracking SDK) in a subsample of 17 participants (due to a technical error no eye tracking data were recorded for the remaining participants). To compensate for the internal processing time of the eye tracker—specifically, the delay between recording an eye tracking sample and the moment when the corresponding sample is provided by the eye tracking SDK (Stein et al., 2021)—and to ensure temporal alignment of the eye tracking data with the EEG data, we first performed a cross-correlation between the eye tracking data and the EOG channels of the EEG data. This yielded a temporal offset between the two signals of on average 60 ms (*SD* = 3.80, range: 50–67 ms); this delay was corrected by subtracting it from the timestamps in the eye tracking samples. We performed a baseline-correction procedure on a trial-by-trial basis, in which we subtracted the mean gaze position during the 200 ms time window before the onset of the stimulus face from the gaze positions in the entire trial. Furthermore, we identified blinks using the algorithm based on the dynamics of the pupil dilation values suggested by Kret & Sjak-Shie (2019), which specifically looks for rapid changes in pupil size. We then linearly interpolated the gaze values during the times of identified blinks (plus a safety margin of 100 ms on each side). We applied the same decoding approach as we used for the EEG data: we trained and evaluated a time-resolved linear classifier with a similar architecture (L2-regularized logistic regression; λ = 1.0; repeated randomized 5-fold cross-validation) for each participant. Here, we decoded the target variables (e.g., facial expressions, depth conditions, face identities) from the time-resolved, two-dimensional gaze direction. To this end, we converted the gaze vector (i.e., viewing direction) into spherical coordinates: phi (vertical) and theta (horizontal component). In contrast to the EEG-decoding, we did not apply downsampling nor mini-batching for the decoding from eye tracking data because, in comparison to EEG, eye tracking data are substantially less prone to measurement noise and single-trial data are more reliable.

In exploratory analyses, we analyzed the gaze behavior by classifying saccades and fixations on a single trial level. Saccade detection was based on a velocity-based algorithm with noise-dependent threshold (i.e., 5 SDs velocity threshold and minimum duration of 3 samples; Engbert et al., 2015; Engbert & Kliegl, 2003; Engbert & Mergenthaler, 2006). For these analyses, we excluded saccades with amplitudes smaller than 2 degrees of visual angle. This exclusion criterion was implemented due to the limitations in precision and sampling rate of the eye tracker within the VR headset, which make the classification and description of smaller saccades unreliable. For fixations, we retained only those with a duration of at least 50 milliseconds. To calculate the fixation heatmaps in Figure 4, we followed the recommendation provided by Le Meur & Baccino (2013): On the subject-level, we calculated the average spherical coordinates for each fixation, binned the fixations into spatial bins of a width and height of 0.1 dva, and convolved the resulting two-dimensional histogram (across all trials of the respective condition) with a two-dimensional gaussian kernel (*SD* = 1 dva for both dimensions and default parameters for the method “gaussian_kernel” from the SciPy Python module). We calculated a 4×2×4 MANOVA which modeled the average fixation position (two-dimensional, spherical coordinates) as a function of the face’s expression (neutral, angry, happy, surprised), depth condition (mono, stereo), and time window (P1, N170, EPN, LPC). We used the same model to analyze the (two-dimensional) saccade directions. Further, we applied Generalized Linear Models (GLM) to model the number of saccades (Poisson, log-linear link function) and the saccade amplitude (Gaussian, log-linear) as a function of emotional expression, time window, and depth condition.

### Behavioral data

We compared participants’ behavioral categorizations of each trial’s facial expression with the objective categorizations based on the FACS values to calculate confusion matrices (Supplementary Figure S1). Due to the high level of agreement, we used the FACS categories as ground truth for the further analyses. To test for differences in the recognition performance between the mono- and the stereoscopic depth condition, we calculated a rmANOVA, modeling the recognition scores (i.e., the values on the diagonal of the confusion matrices in Supplementary Figure S1) as a function of the depth condition (two levels), the emotional expression (four levels), and their interaction. Finally, we modeled the intensity ratings from the last part of the experiment (based on the 9-point SAM scale) using a rmANOVA with the factors emotional expression and depth condition.

## Supplementary Material

Supplementary information is available in a separate file.

## Data availability

The data that support the findings of this study are openly available in “Edmond – the Open Research Data Repository of the Max Planck Society” at https://doi.org/10.17617/3.CQ2VXX. The code for data analysis can be found at https://github.com/eioe/vr2f.

## Acknowledgements

We thank Michael Gilbert for his support and guidance regarding the stimulus material, particularly for providing the original 3D face models from his validation study. We are also grateful to Jeroen de Mooij for his assistance and expertise in implementing the experiment in Unity. This research was supported by the cooperation between the Max Planck Society and the Fraunhofer Gesellschaft (grant: project NEUROHUM), by the German Federal Ministry of Education and Research (BMBF/BMFTR) under grants 13GW0206, 13GW0488, 16SV9156, and by the Deutsche Forschungsgemeinschaft (DFG) under grants 502864329 and 542559580. Relevant parts of this work were conducted at the Max Planck Dahlem Campus of Cognition (MPDCC) of the Max Planck Institute for Human Development, Berlin, Germany.

## CrediT statement

**Felix Klotzsche:** Conceptualization, Data curation, Formal Analysis, Investigation, Methodology, Software, Visualization, Writing – original draft, Writing – review & editing

**Ammara Nasim:** Conceptualization, Data curation, Investigation, Methodology, Writing – review & editing;

**Simon M. Hofmann:** Conceptualization, Methodology, Software, Writing – review & editing;

**Arno Villringer:** Funding Acquisition, Resources, Supervision;

**Vadim Nikulin:** Formal Analysis, Methodology, Supervision, Writing – review & editing;

**Werner Sommer:** Conceptualization, Formal Analysis, Methodology, Supervision, Visualization, Writing – review & editing;

**Michael Gaebler:** Conceptualization, Funding Acquisition, Resources, Supervision, Writing – original draft, Writing – review & editing;

## Attributions

Figure 1:

*Canon AT-1 Retro Camera* (https://skfb.ly/6ZwNB) by AleixoAlonso and *EyeBall* (https://skfb.ly/osJMS) by CrisLArt are licensed under Creative Commons Attribution (http://creativecommons.org/licenses/by/4.0/).

## Footnotes

1 Successfully decoding a stimulus feature suggests that the decodable information is present in the recorded data and—given some necessary assumptions about their origin—also available in the identified physiological sources (e.g., brain regions at a given moment in time). This does not imply that the brain (e.g., other brain regions) actually makes use of this information—which would be a necessary requirement for a representation in a more strict interpretation (Kriegeskorte & Douglas, 2019; Ritchie et al., 2019).

