## Supplementary Material for "EEG decodability of facial expressions and their stereoscopic depth cues in immersive virtual reality"

### Figures

#### S1: Behavioral confusion matrices

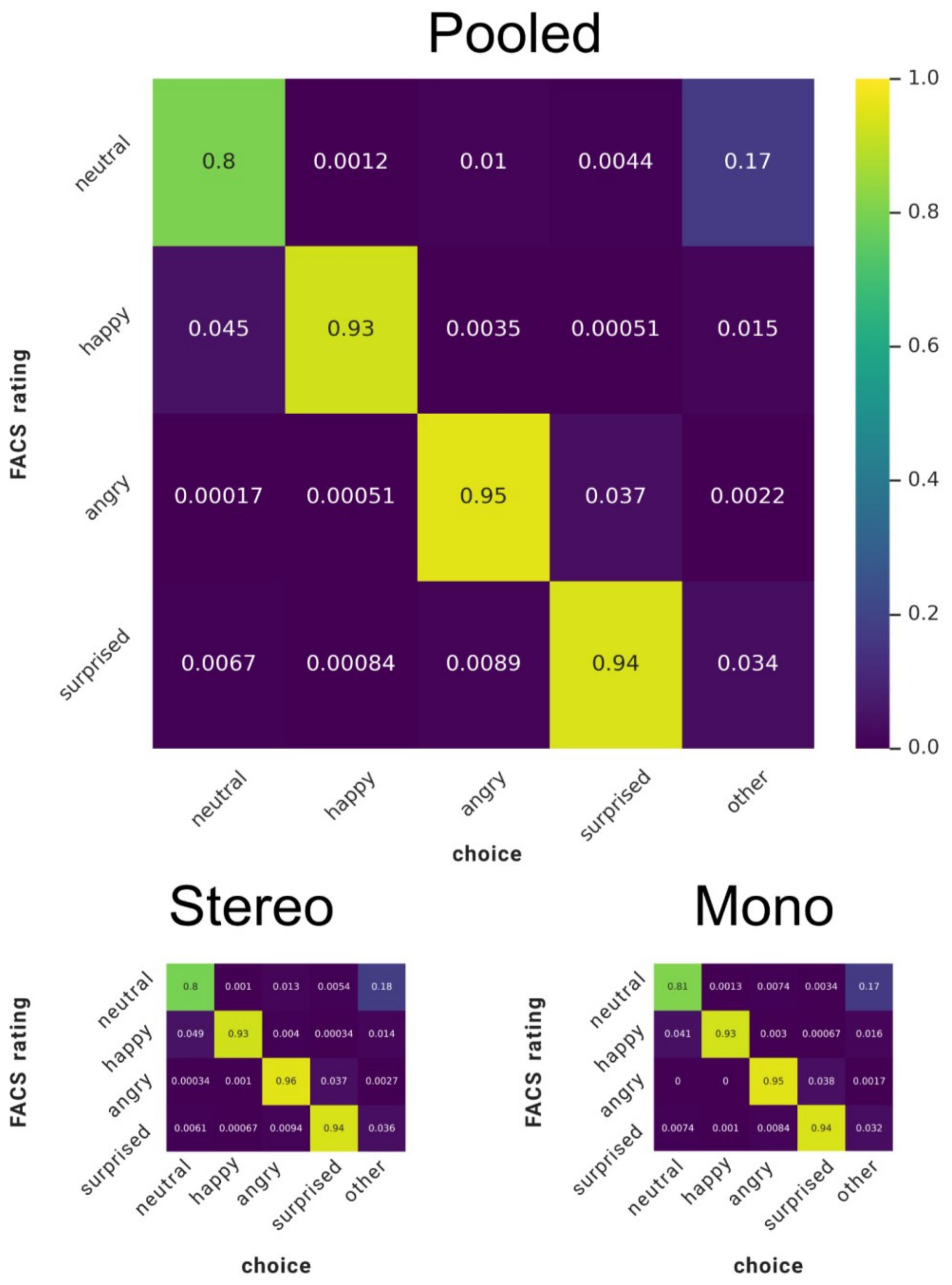

**Figure S1:** Behavioral confusion matrices. Results are shown pooled across viewing conditions (top) and separately for conditions with stereoscopic depth cues (*Stereo*) and without them (*Mono*) (bottom). *FACS rating* refers to the facial expression label based on the Facial Action Coding System; *choice* refers to the label assigned by the participant. The numbers in the diagonal cells indicate the level of agreement.

#### S2: Intensity ratings

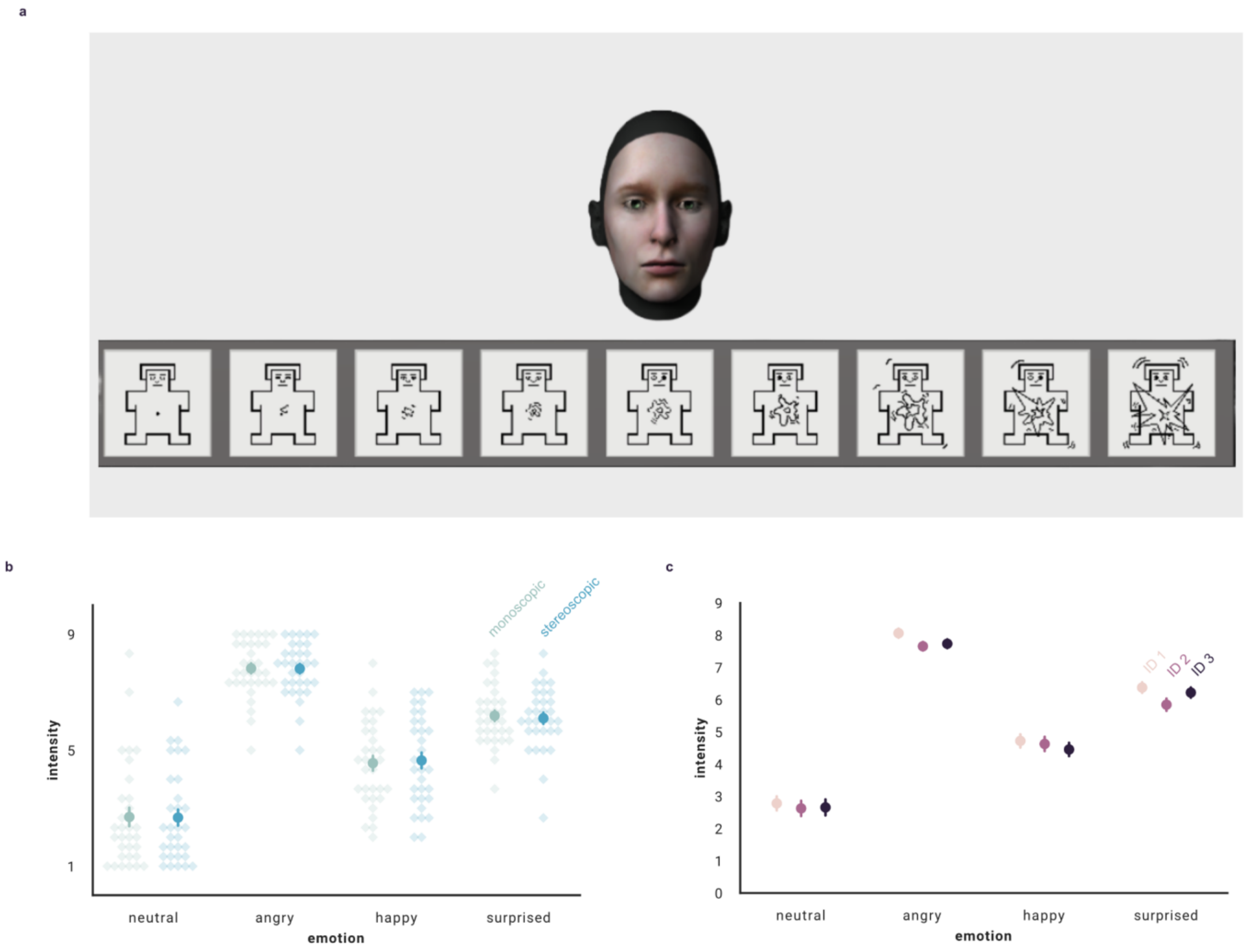

**Figure S2:** At the end of the experiment, participants rated the intensity of the expressed emotion for each of the stimuli. (a) Example trial: a 9-value SAM rating scale (Bradley & Lang, 1994) was shown below the stimulus face. Participants used the VR controller to choose the manikin corresponding to their assessment. (b) Intensity ratings as a function of the displayed facial expression(x-axis) and the viewing condition (color). (c) Link in (b) but for the three different stimulus identities.

#### S3: Pairwise-decoding contrasts (emotional expressions)

**(a)**

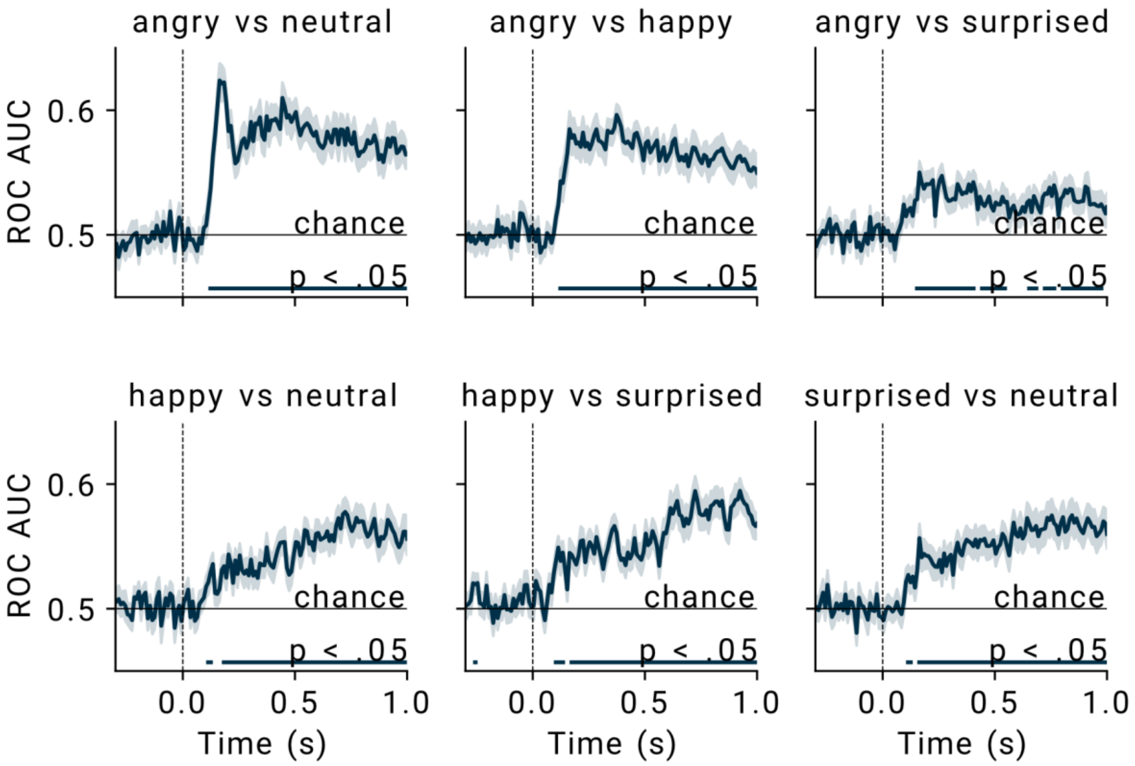

**b)**

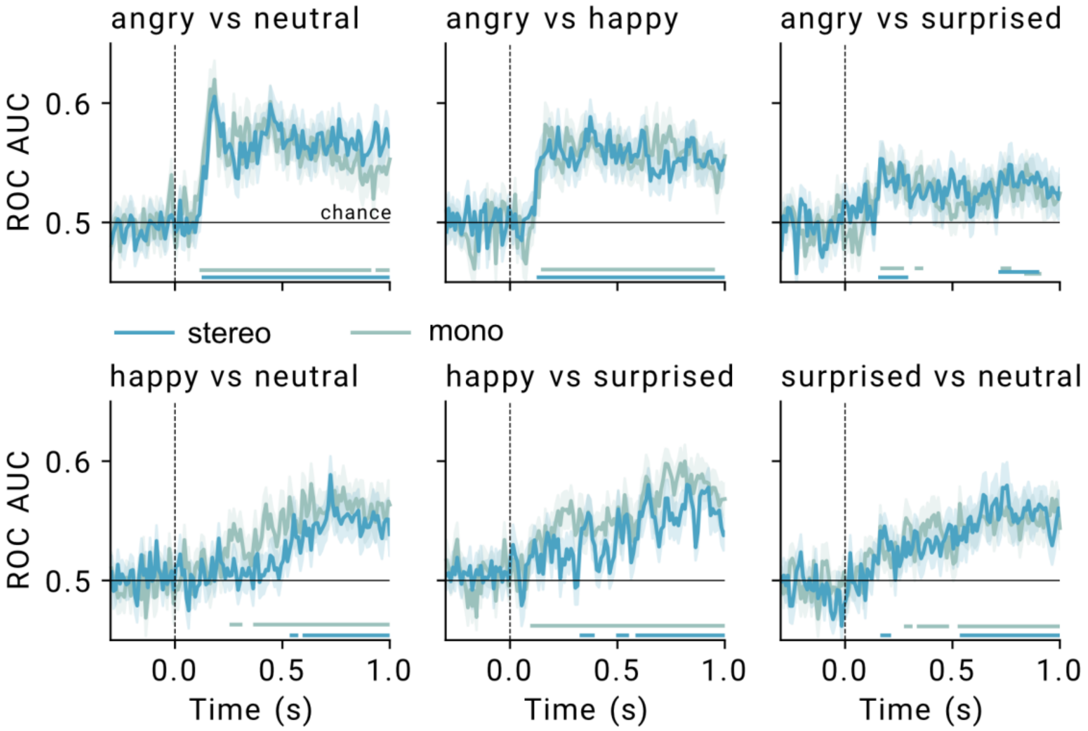

**Figure S3:** Decoding performance over time for the six binary contrasts, comparing pairs of emotions. **(a)** Across both viewing conditions. **(b)** Separately for monoscopic and stereoscopic trials.

#### S4: Pairwise contrasts (emotional expressions) per time window

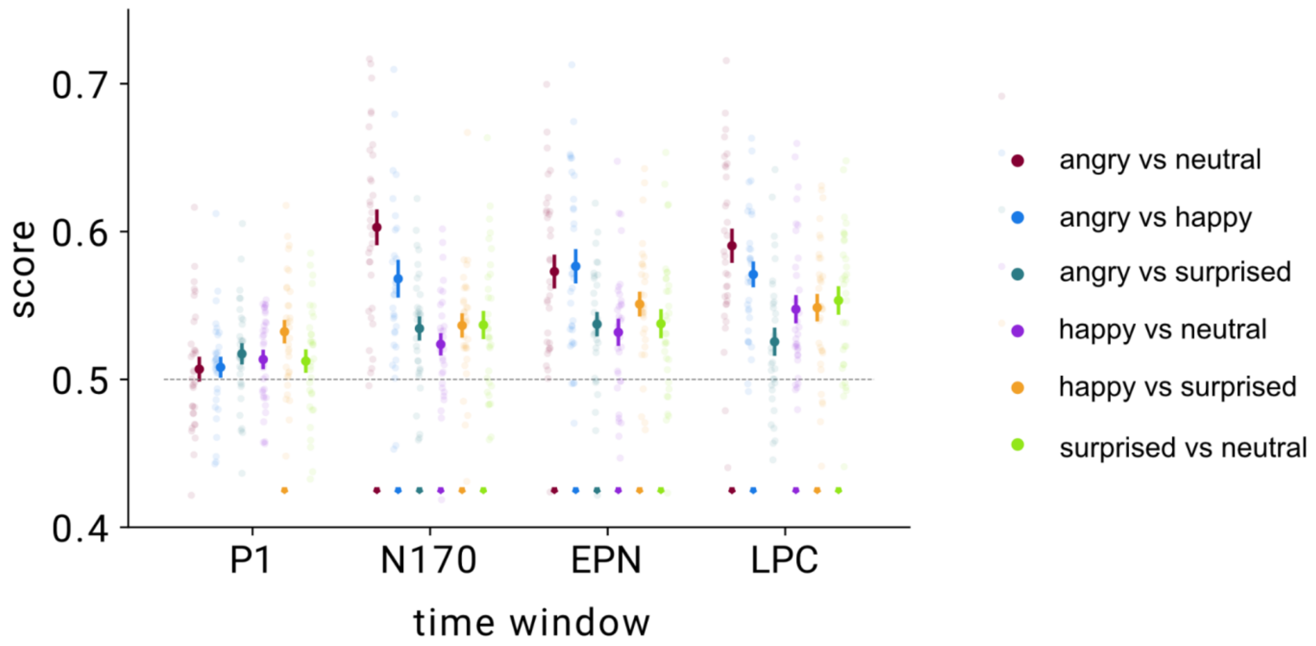

**Figure S4:** Average decoding performance (ROC-AUC) per time window and binary contrast.

#### S5: Cross-decoding results: train on monoscopic → test on stereoscopic and v.v. (decoding target: facial expression)

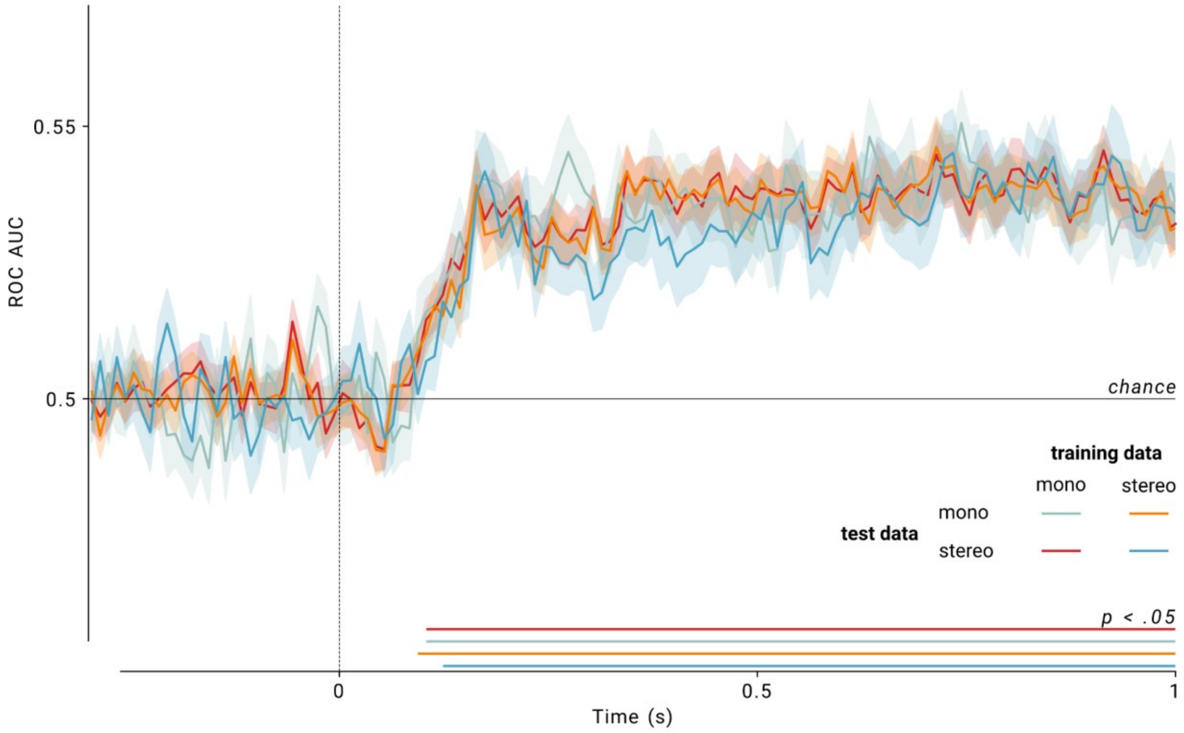

**Figure S5:** Cross-decoding results. Time-resolved decoding performance for classifiers trained to distinguish between the four facial expressions. Separate classifiers were trained on data from the mono- and stereoscopic viewing conditions. Each classifier was then tested either on left-out trials from the same viewing condition (blueish lines) or on trials from the other viewing condition (orange and red lines). Decoding performance did not differ between within- and across-condition tests (no significant clusters when testing for the difference).

#### S6: Fixation heatmaps per time window

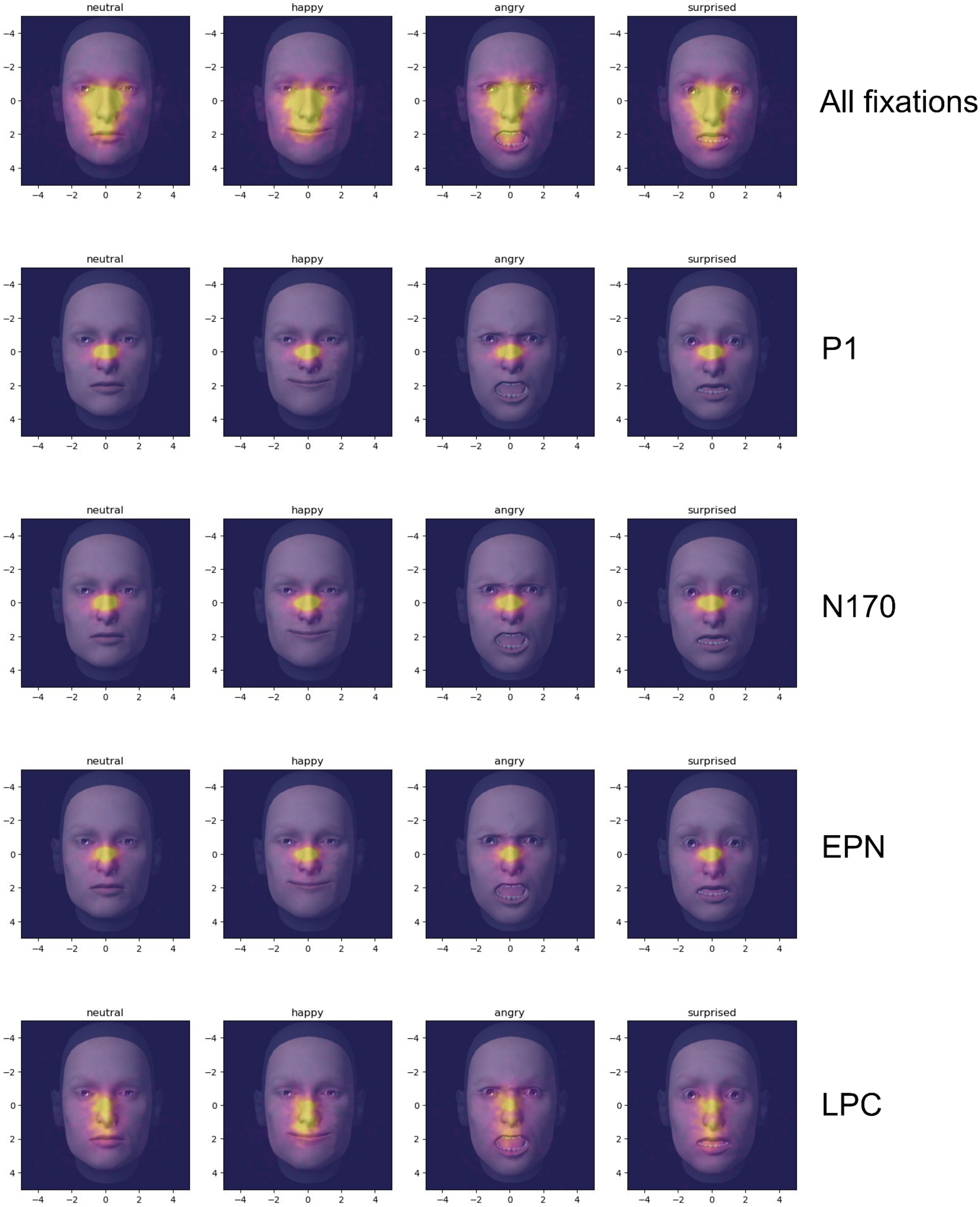

#### S7: Correlation between EEG and eye tracking decoding performances

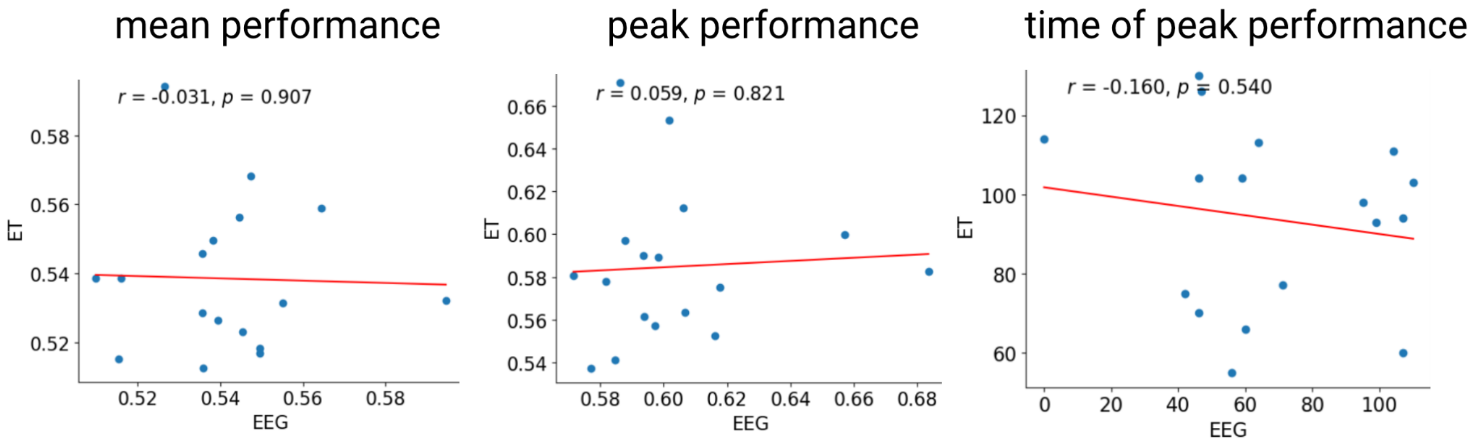

**Figure S7:** No correlations between decoding performance metrics of the classifiers (decoding target: facial expression) trained on EEG and eye tracking (ET) data. Data points are single participants. (N=17, since we did not have eye tracking data for all participants.)

#### S8: Full source reconstruction results

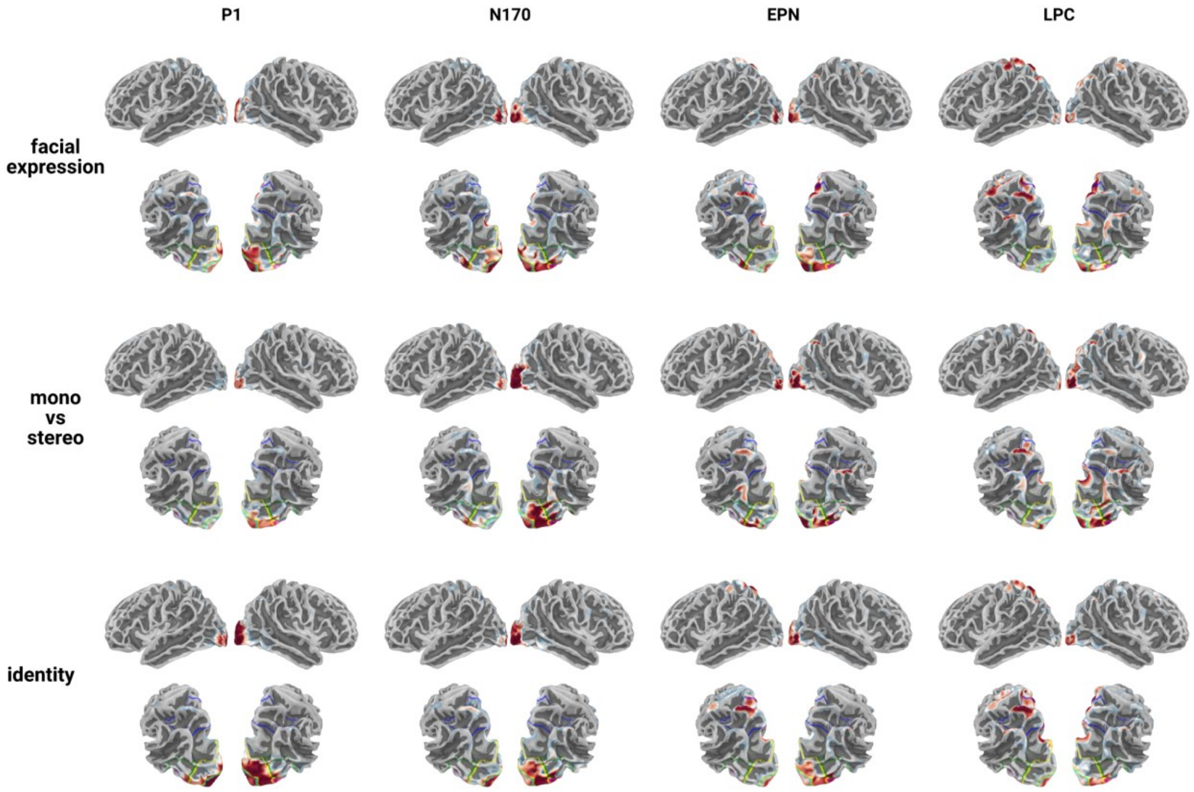

**Figure S8:** Comparison of the reconstruction results for the most informative cortical sources for the decoders distinguishing between the facial expressions (top row), trials with and without stereoscopic depth information (middle row), and between the three stimulus identities (bottom row). Each row shows lateral views (upper half) and parietal views (lower half) on the two hemispheres. Warmer colors indicate more informative sources. The colored lines on the parietal views mark the borders of relevant cortical regions (for details see Figure 3).

#### S9: Source reconstruction results for binary classifiers

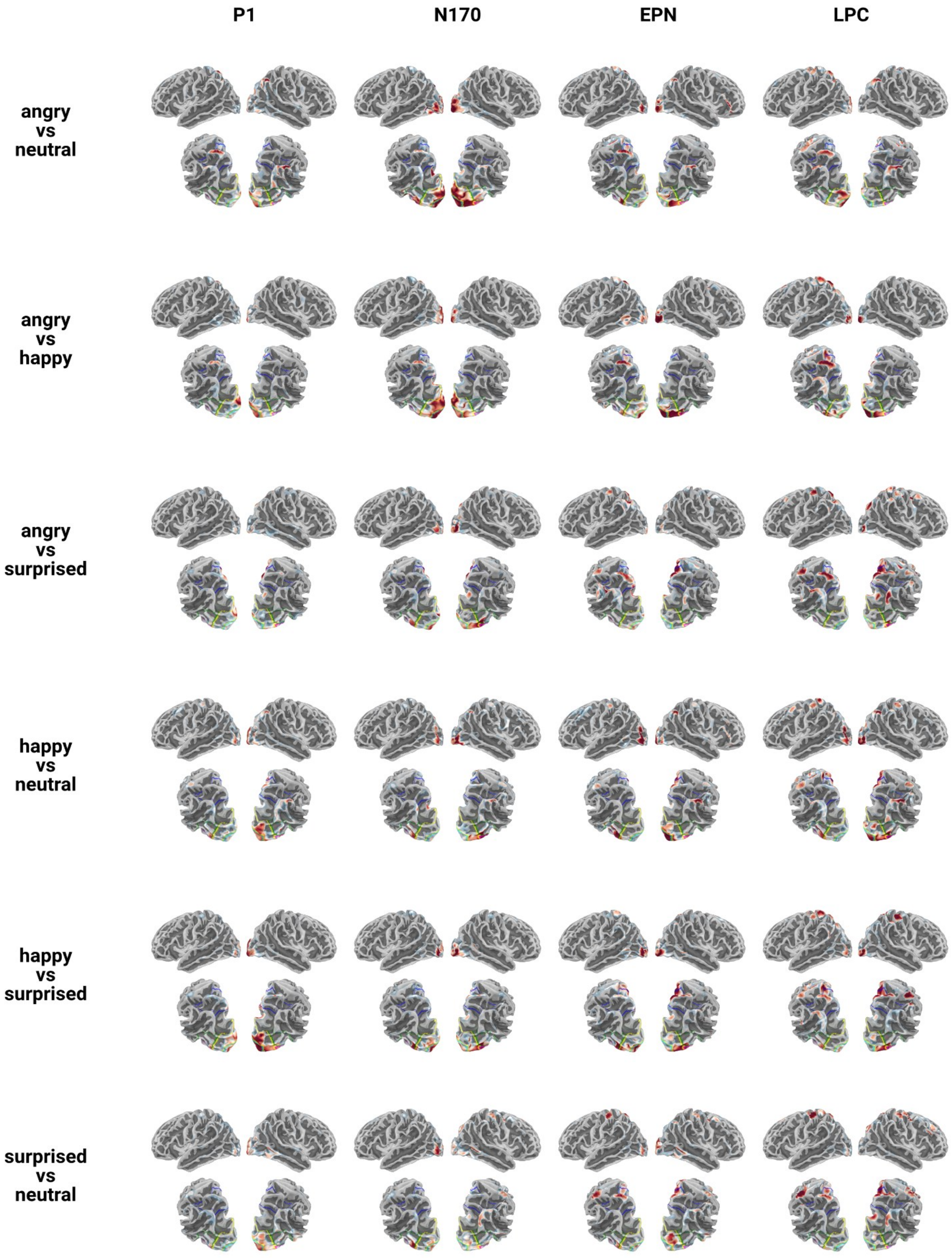

**Figure S9:** Source reconstruction results for the binary classifiers, separately for the four time windows. For details see Figures 3 and S8.

### Tables

#### ST1: Peak EEG-Decoding per Contrast and Viewing Condition

**Table ST1:** Peak EEG-decoding score (1) and time (2) for the binary decoders comparing pairs of emotions, as well as the classifier distinguishing the three different stimulus identities (task-irrelevant feature). Separately for the two viewing conditions. Last columns: Condition comparison results in the form of a paired *t*-test. *Mono*: monoscopic condition; *Stereo*: stereoscopic condition.

##### 1 Peak decoding score

| | Mono | | | Stereo | | | Stats | |
|---|---|---|---|---|---|---|---|---|
| | Mean | SD | CI | Mean | SD | CI | t(32) | p |
| angry vs neutral | 0.74 | 0.06 | [0.72, 0.76] | 0.74 | 0.05 | [0.72, 0.76] | 0.17 | 0.869 |
| angry vs happy | 0.73 | 0.05 | [0.71, 0.74] | 0.72 | 0.06 | [0.70, 0.74] | 0.42 | 0.679 |
| angry vs surprised | 0.70 | 0.04 | [0.69, 0.72] | 0.70 | 0.05 | [0.68, 0.71] | 0.65 | 0.523 |
| happy vs neutral | 0.72 | 0.04 | [0.70, 0.73] | 0.70 | 0.05 | [0.69, 0.72] | 1.98 | 0.056 |
| happy vs surprised | 0.72 | 0.04 | [0.71, 0.74] | 0.71 | 0.05 | [0.69, 0.73] | 1.38 | 0.176 |
| surprised vs neutral | 0.72 | 0.05 | [0.70, 0.74] | 0.72 | 0.06 | [0.70, 0.74] | 0.23 | 0.821 |
| id1 vs id2 vs id3 | 0.60 | 0.02 | [0.60, 0.61] | 0.61 | 0.03 | [0.60, 0.62] | -1.44 | 0.159 |

##### 2 Time of peak decoding score

| | Mono | | | Stereo | | | Stats | |
|---|---|---|---|---|---|---|---|---|
| | Median | SD | CI | Median | SD | CI | t(32) | p |
| angry vs neutral | 344.00 | 230.94 | [265.20, 422.80] | 454.00 | 363.03 | [330.14, 577.86] | -0.89 | 0.381 |
| angry vs happy | 334.00 | 278.11 | [239.11, 428.89] | 344.00 | 307.00 | [239.25, 448.75] | -0.59 | 0.561 |
| angry vs surprised | 184.00 | 311.88 | [77.59, 290.41] | 194.00 | 272.23 | [101.12, 286.88] | -0.74 | 0.462 |
| happy vs neutral | 294.00 | 370.40 | [167.62, 420.38] | 574.00 | 402.62 | [436.63, 711.37] | -1.70 | 0.100 |
| happy vs surprised | 624.00 | 384.09 | [492.95, 755.05] | 364.00 | 391.24 | [230.51, 497.49] | 1.36 | 0.184 |
| surprised vs neutral | 404.00 | 357.74 | [281.94, 526.06] | 544.00 | 376.77 | [415.45, 672.55] | -0.31 | 0.756 |
| id1 vs id2 vs id3 | 124.00 | 293.89 | [23.73, 224.27] | 94.00 | 298.97 | [-8.01, 196.01] | 1.01 | 0.320 |

### ST2: Most informative source per time window and contrast

**Table ST2:** Most informative cortical region per decoding target and time window. Regions as defined by the reduced parcellation atlas (23 labels per hemisphere) provided by Glasser et al. (2016) and Mills (2016).

#### A) Both depth conditions

Most informative sources (both viewing conditions)

| time win | start-end | mono vs stereo | neutral vs happy vs angry vs surprised | id1 vs id2 vs id3 |
|---|---|---|---|---|
| P1 | 0.08-0.12 | Primary Visual Cortex (V1)-rh | Early Visual Cortex-rh | Primary Visual Cortex (V1)-lh |
| N170 | 0.13-0.2 | Ventral Stream Visual Cortex-rh | Early Visual Cortex-rh | Ventral Stream Visual Cortex-rh |
| EPN | 0.25-0.3 | MT+ Complex and Neighboring Visual Areas-rh | Primary Visual Cortex (V1)-rh | Superior Parietal Cortex-lh |
| LPC | 0.4-0.6 | Ventral Stream Visual Cortex-rh | Superior Parietal Cortex-rh | Superior Parietal Cortex-rh |

| time win | start-end | angry vs neutral | angry vs happy | angry vs surprised |
|---|---|---|---|---|
| P1 | 0.08-0.12 | Inferior Parietal Cortex-rh | Early Visual Cortex-rh | Superior Parietal Cortex-rh |
| N170 | 0.13-0.2 | Early Visual Cortex-rh | Early Visual Cortex-rh | Ventral Stream Visual Cortex-rh |
| EPN | 0.25-0.3 | Early Visual Cortex-rh | Primary Visual Cortex (V1)-rh | Somatosensory and Motor Cortex-rh |
| LPC | 0.4-0.6 | Primary Visual Cortex (V1)-lh | Primary Visual Cortex (V1)-rh | Premotor Cortex-rh |

| time win | start-end | happy vs neutral | happy vs surprised | surprised vs neutral |
|---|---|---|---|---|
| P1 | 0.08-0.12 | Primary Visual Cortex (V1)-lh | Early Visual Cortex-rh | Primary Visual Cortex (V1)-rh |
| N170 | 0.13-0.2 | Ventral Stream Visual Cortex-rh | MT+ Complex and Neighboring Visual Areas-lh | MT+ Complex and Neighboring Visual Areas-lh |
| EPN | 0.25-0.3 | Primary Visual Cortex (V1)-rh | Early Visual Cortex-lh | Somatosensory and Motor Cortex-lh |
| LPC | 0.4-0.6 | Superior Parietal Cortex-rh | Superior Parietal Cortex-rh | Somatosensory and Motor Cortex-lh |

#### B) Monoscopic depth condition

Most informative sources (monoscopic condition)

| time win | start-end | neutral vs happy vs angry vs surprised | id1 vs id2 vs id3 |
|---|---|---|---|
| P1 | 0.08-0.12 | Primary Visual Cortex (V1)-rh | Early Visual Cortex-rh |
| N170 | 0.13-0.2 | Ventral Stream Visual Cortex-rh | Ventral Stream Visual Cortex-rh |
| EPN | 0.25-0.3 | Early Visual Cortex-lh | Superior Parietal Cortex-lh |
| LPC | 0.4-0.6 | Dorso-Lateral Prefrontal Cortex-lh | Superior Parietal Cortex-lh |

| time win | start-end | angry vs neutral | angry vs happy | angry vs surprised |
|---|---|---|---|---|
| P1 | 0.08-0.12 | Superior Parietal Cortex-lh | Primary Visual Cortex (V1)-lh | DorsoLateral Prefrontal Cortex-lh |
| N170 | 0.13-0.2 | Early Visual Cortex-lh | Primary Visual Cortex (V1)-lh | Ventral Stream Visual Cortex-rh |
| EPN | 0.25-0.3 | Superior Parietal Cortex-rh | Early Visual Cortex-lh | Ventral Stream Visual Cortex-rh |
| LPC | 0.4-0.6 | Superior Parietal Cortex-rh | DorsoLateral Prefrontal Cortex-lh | DorsoLateral Prefrontal Cortex-lh |

| time win | start-end | happy vs neutral | happy vs surprised | surprised vs neutral |
|---|---|---|---|---|
| P1 | 0.08-0.12 | Superior Parietal Cortex-rh | MT+ Complex and Neighboring Visual Areas-lh | Somatosensory and Motor Cortex-lh |
| N170 | 0.13-0.2 | Superior Parietal Cortex-rh | Primary Visual Cortex (V1)-lh | Somatosensory and Motor Cortex-lh |
| EPN | 0.25-0.3 | Paracentral Lobular and Mid Cingulate Cortex-lh | MT+ Complex and Neighboring Visual Areas-lh | Somatosensory and Motor Cortex-lh |
| LPC | 0.4-0.6 | Superior Parietal Cortex-rh | Paracentral Lobular and Mid Cingulate Cortex-lh | Somatosensory and Motor Cortex-lh |

### C) Stereoscopic depth condition

#### Most informative sources (stereoscopic condition)

| time win | start-end | neutral vs happy vs angry vs surprised | id1 vs id2 vs id3 |
|---|---|---|---|
| P1 | 0.08-0.12 | Primary Visual Cortex (V1)-rh | Primary Visual Cortex (V1)-rh |
| N170 | 0.13-0.2 | Early Visual Cortex-rh | MT+ Complex and Neighboring Visual Areas-rh |
| EPN | 0.25-0.3 | Superior Parietal Cortex-lh | Primary Visual Cortex (V1)-rh |
| LPC | 0.4-0.6 | Superior Parietal Cortex-rh | Somatosensory and Motor Cortex-rh |

| time win | start-end | angry vs neutral | angry vs happy | angry vs surprised |
|---|---|---|---|---|
| P1 | 0.08-0.12 | Primary Visual Cortex (V1)-rh | Early Visual Cortex-lh | Lateral Temporal Cortex-rh |
| N170 | 0.13-0.2 | Dorsal Stream Visual Cortex-lh | Early Visual Cortex-lh | MT+ Complex and Neighboring Visual Areas-lh |
| EPN | 0.25-0.3 | Superior Parietal Cortex-lh | Superior Parietal Cortex-lh | Superior Parietal Cortex-lh |
| LPC | 0.4-0.6 | Superior Parietal Cortex-lh | Superior Parietal Cortex-rh | Paracentral Lobular and Mid Cingulate Cortex-rh |

| time win | start-end | happy vs neutral | happy vs surprised | surprised vs neutral |
|---|---|---|---|---|
| P1 | 0.08-0.12 | Early Visual Cortex-rh | Primary Visual Cortex (V1)-rh | Dorsal Stream Visual Cortex-rh |
| N170 | 0.13-0.2 | Dorsal Stream Visual Cortex-lh | Superior Parietal Cortex-rh | Somatosensory and Motor Cortex-rh |
| EPN | 0.25-0.3 | Primary Visual Cortex (V1)-rh | Superior Parietal Cortex-lh | Somatosensory and Motor Cortex-lh |
| LPC | 0.4-0.6 | Primary Visual Cortex (V1)-rh | Superior Parietal Cortex-rh | Dorsal Stream Visual Cortex-rh |

#### ST3: Eye Tracking: Number of saccades as function of emotion * viewing condition * time window (GLM)

We fitted a Poisson GLM with a log link function to model the number of saccades (`sacc_count`) as a function of emotion (emotion), viewing condition (viewcond), and EEG time window (tw), including all two- and three-way interactions. Because the time windows varied in length, we included the logarithm of the time window duration as an offset term to model saccade rate (i.e., saccades per unit time).

The model was specified as follows:

```
glm("sacc_count ~ C(emotion, Treatment('neutral')) *
                      C(viewcond, Treatment('mono')) *
                       C(tw, Treatment('p1'))",
    data=saccades_count_tw,
    family=sm.families.Poisson(),
    offset=np.log(saccades_count_tw["window_length"]))
```

**Results:**

```
Generalized Linear Model Regression Results
Dep. Variable:      sacc_count   No. Observations:          517
Model:                     GLM   Df Residuals:              485
Model Family:          Poisson   Df Model:                   31
Link Function:             Log   Scale:                  1.0000
Method:                   IRLS   Log-Likelihood:        -2763.7
Deviance:               3333.1
Pearson chi2:         3.68e+03
Pseudo R-squ. (CS):     0.8967
Covariance Type:     nonrobust
```

| Factor | chi2 | P>chi2 | df constraint |
|---|---|---|---|
| Intercept | 1948.99948 | 0.000000e+0 | 1 |
| Emotion | 2.415737 | 4.907120e-01 | 3 |
| Viewcond | 1.473101 | 2.248571e-01 | 1 |
| Time window | 115.813726 | 6.150233e-25 | 3 |
| Emotion:Viewcond | 1.050514 | 7.890316e-01 | 3 |
| Emotion:Time window | 4.803167 | 8.511180e-01 | 9 |
| Viewcond:Time window | 3.979863 | 2.636468e-01 | 3 |
| Emotion:Viewcond:Time window | 3.082237 | 9.609450e-01 | 9 |

| Predictor | β (SE) | z | p | 95% CI |
|---|---|---|---|---|
| Intercept | 4.91 (0.11) | 44.15 | < .001 | [4.69, 5.12] |
| tw[EPN] | 0.86 (0.13) | 6.75 | < .001 | [0.61, 1.10] |
| tw[LPC] | 0.19 (0.12) | 1.57 | 0.117 | [–0.05, 0.42] |
| tw[N170] | –0.07 (0.14) | –0.51 | 0.614 | [–0.35, 0.21] |
| emotion × viewcond × tw | All p > .05 | — | — | |
